# A novel subgroup of light-driven sodium pumps with an additional Schiff base counterion

**DOI:** 10.1101/2023.10.11.561842

**Authors:** E. Podoliak, G. H. U. Lamm, A. Alekseev, E. Marin, A. V. Schellbach, D. A. Fedotov, A. Stetsenko, N. Maliar, G. Bourenkov, T. Balandin, C. Baeken, R. Astashkin, T. R. Schneider, A. Bateman, J. Wachtveitl, I. Schapiro, V. Busskamp, A. Guskov, V. Gordeliy, K. Kovalev

**Affiliations:** Department of Ophthalmology, University Hospital Bonn, Medical Faculty, Bonn, Germany; Institute of Physical and Theoretical Chemistry, Goethe University Frankfurt, 60438 Frankfurt am Main, Germany; Advanced Optogenes Group, Institute for Auditory Neuroscience and InnerEarLab, University Medical Center Göttingen, Göttingen, Germany; Cluster of Excellence “Multiscale Bioimaging: from Molecular Machines to Networks of Excitable Cells” (MBExC), University of Göttingen, Göttingen, Germany; Groningen Institute for Biomolecular Sciences and Biotechnology, University of Groningen, 9747AG, Groningen, the Netherlands; Fritz Haber Center for Molecular Dynamics Research, Institute of Chemistry, The Hebrew University of Jerusalem, Jerusalem, 9190401, Israel; Department of Biochemistry, University of Cambridge, 80 Tennis Court Road, Cambridge CB2 1GA, UK; European Molecular Biology Laboratory, EMBL Hamburg c/o DESY, 22607, Hamburg, Germany; Institute of Biological Information Processing (IBI-7: Structural Biochemistry), Forschungszentrum Jülich, Jülich, Germany; JuStruct: Jülich Center for Structural Biology, Forschungszentrum Jülich, Jülich, Germany; Univ. Grenoble Alpes, CEA, CNRS, Institut de Biologie Structurale (IBS), 38000, Grenoble, France; European Molecular Biology Laboratory, European Bioinformatics Institute (EMBL-EBI), Wellcome Genome Campus, Hinxton, UK

## Abstract

Light-driven sodium pumps (NaRs) are unique ion-transporting microbial rhodopsins. The major group of NaRs is characterized by an NDQ motif and has two aspartic acid residues in the central region essential for sodium transport. Here we identified a new subgroup of the NDQ rhodopsins bearing an additional glutamic acid residue in the close vicinity to the retinal Schiff base. We thoroughly characterized a member of this subgroup, namely the protein *Er*NaR from *Erythrobacter sp. HL-111* and showed that the additional glutamic acid results in almost complete loss of pH sensitivity for sodium-pumping activity, which is in contrast to previously studied NaRs. *Er*NaR is capable of transporting sodium efficiently even at acidic pH levels. X-ray crystallography and single particle cryo-electron microscopy reveal that the additional glutamic acid residue mediates the connection between the other two Schiff base counterions and strongly interacts with the aspartic acid of the characteristic NDQ motif. Hence, it reduces its pKa. Our findings shed light on a new subgroup of NaRs and might serve as a basis for their rational optimization for optogenetics.

## MAIN

Light-driven sodium pumps (NaRs) were discovered in 2013 with the characterization of the microbial rhodopsin (MR) KR2 from the bacterium *Krokinobacter eikastus*^1^. NaRs are membrane proteins that actively transport sodium outside of the cell in response to light illumination^2^. Since 2013, numerous various NaRs have been identified^3–6^. Most of them have an NDQ motif (N112, D116, and Q123 residues in KR2 corresponding to D85, T89, and D96 residues of the proton pump bacteriorhodopsin (BR^7^)) in the helix C. Another group of MRs able to pump sodium was recently reported to have a DTG motif^8^.

As all other MRs, the structure of NDQ rhodopsins consists of seven transmembrane helices (A-G) encapsulating a retinal cofactor, covalently attached to the lysine residue of the helix G (K255 in KR2) via a Schiff base (RSB)^1,9,10^. NDQ rhodopsins form pentamers in the membrane as was shown for KR2^9,11,12^. The transported substrate, sodium, is not bound inside these rhodopsins in the non-illuminated (resting) state, but was found at the oligomerization interface of KR2 coordinated by two neighboring protomers^9,12^. Upon light illumination, NDQ rhodopsins undergo a photocycle with several intermediate states, transitions between which result in sodium translocation across the membrane. Namely, there are the K, L, M, and O intermediates of a typical NDQ rhodopsin, where the O state was shown to be the only one associated with the transient sodium binding inside the protein^1^. For several NDQ rhodopsins, such as KR2 and the NaR from *Indibacter alkaliphilus* (*Ia*NaR), the O state was demonstrated to consist of several sub-states^13–18^.

There is still no complete understanding of the molecular mechanism of light-driven sodium pumping by NDQ rhodopsins despite the thorough characterization of KR2. For the latter it was shown that the conformational change in the RSB region takes place upon sodium binding in the O state^14,19^. N112 of the NDQ motif, which points outside of the protomer towards the pentamerization interface in the ground state, flips inside the protein to coordinate the transiently-bound sodium ion^12,19^. This considerable movement (∼5 Å in amplitude) is accompanied by the flip of the L74 side chain, creating sufficient space for N112 inside the KR2 protomer^19^. Such a synchronous switch of the N112-L74 pair is believed to be one of the key determinants of light-driven sodium pumping^19^. Surprisingly, while the role of N112 of that pair in the KR2 functioning was studied in detail^20,21^, the L74 residue often remained out of the focus despite its involvement in sodium-pumping-associated conformational changes. Nevertheless, it was evidenced by the L74A mutation, which dramatically decreases the sodium-pumping activity of KR2, suggesting its important role in the NDQ rhodopsin^19^.

Here, we bioinformatically analyzed the clade of NDQ rhodopsins and found that although the majority of the proteins possess leucine/isoleucine at the position of L74 in KR2, there is a subgroup of proteins bearing glutamate at this position. To see what effect it has on protein structure and function we investigated a rhodopsin from *Erythrobacter sp. HL-111*, *Er*NaR (Uniprot ID: A0A1H1XA63), belonging to this newly identified subgroup. *Er*NaR, in contrast to KR2, demonstrated unique spectroscopic properties with only a minor dependence of absorption spectra on pH. Furthermore, *Er*NaR is efficient in pumping sodium with high selectivity over protons even at acidic pH values as low as 5.0. Single particle cryo-electron microscopy (cryo-EM) and X-ray crystallography showed an unusual conformation of the rhodopsin in the resting state with a very short H-bond between E64 and D105 (corresponding to L74 and D116 in KR2, respectively). These findings provide essential information on the mechanisms of NDQ rhodopsins and natural ways of tuning of their functional properties.

## RESULTS AND DISCUSSION

### Two subgroups of NaRs

Initially, we performed a bioinformatic analysis of the rhodopsins possessing the NDQ motif in available gene databases (UniProtKB, UniParc, GenBank, and MGnify) to obtain the complete list of members of the clade. In total, we identified 351 unique complete sequences of NDQ rhodopsins which resulted in 219 sequences with less than 90% pairwise sequence identity. The proteins cluster into several branches in the phylogenetic tree (Fig. 1A). Closer analysis of these branches showed that, while there are some interesting features of their representatives, the overall clade of NDQ rhodopsins can be divided into two major subgroups, Subgroup 1 and Subgroup 2 (Fig. 1A). Members of the Subgroup 1 possess leucine or isoleucine at the position of L74 in KR2 (Fig. 1B). KR2 belongs to Subgroup 1, which is also the larger one. Members of Subgroup 2 possess glutamic acid at the position of L74 in KR2 (Fig. 1B). Importantly, this position is located close to the RSB and L74 is involved in conformational changes in KR2 associated with sodium translocation^12,19^ (Fig. 1C). Thus, the introduction of glutamic acid might significantly affect the functional, spectroscopic, and structural properties of the NDQ rhodopsins belonging to Subgroup 2. In addition, although the internal polar and rechargeable residues are conserved within the entire NDQ rhodopsins clade (Fig. 1B, left), the amino acid residues at the surface, including those of the oligomerization interface, are significantly different between the two subgroups (Fig. 1B, right). For instance, the D102 residue, forming the interprotomeric sodium binding site on the extracellular surface of the KR2 pentamer^9^, is either absent or substituted with glutamic acid in the Subgroup 2 (Fig. 1B,D).

**Fig. 1.**
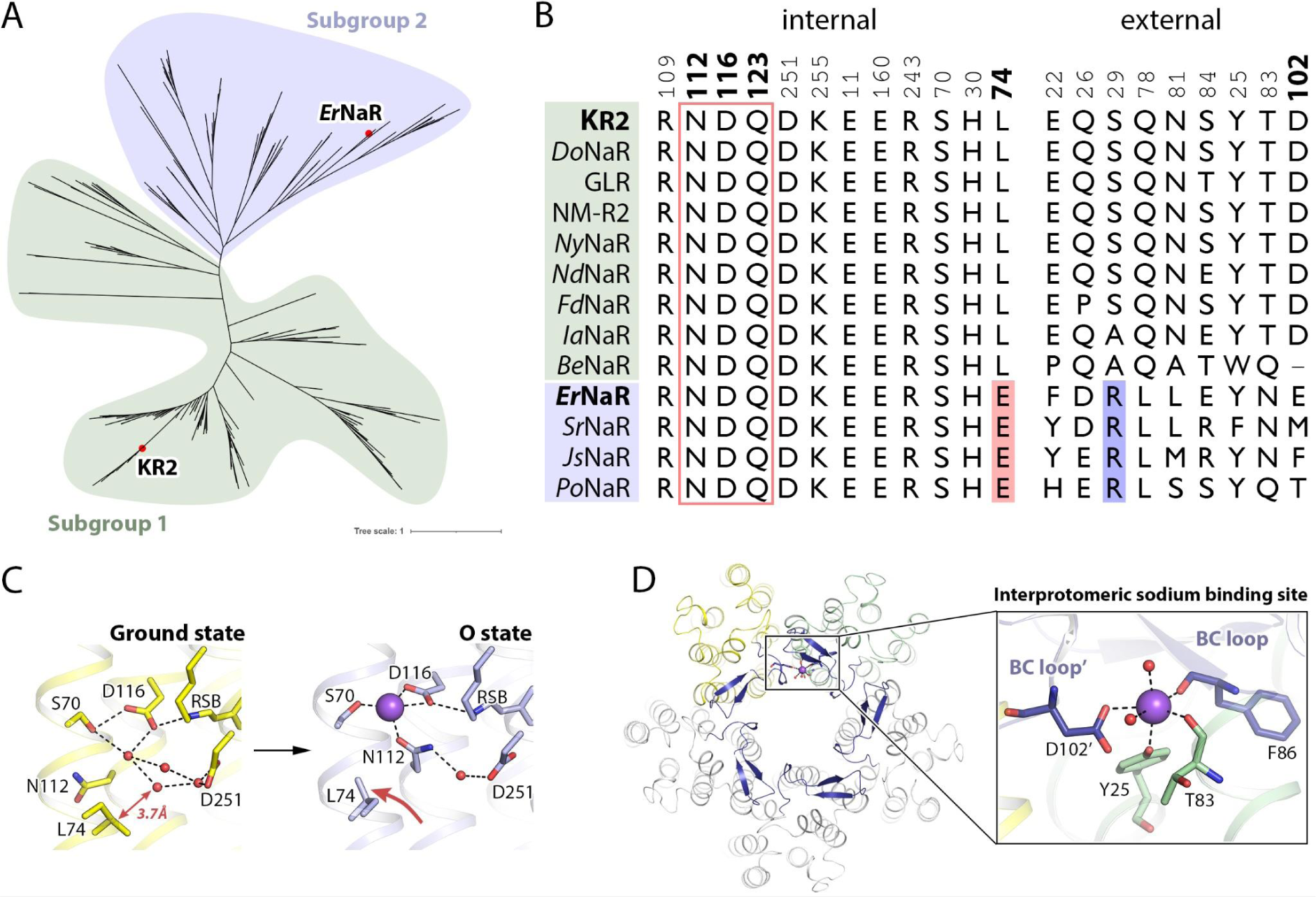
Phylogeny of NDQ rhodopsins. **A.** Phylogenetic tree of the NDQ rhodopsins clade. Subgroups 1 and 2 are highlighted in green and blue, respectively. **B.** Sequence alignment of the key regions of the representative NDQ rhodopsins: internal polar/rechargeable amino acid residues (left), functionally and structurally relevant external amino acid residues (right). The key positions are marked with bold numbers according to the KR2 sequence. Additional RSB counterion (E64 in *Er*NaR) in the Subgroup 2 is highlighted red. Additional positively charged residue near the interprotomeric sodium binding site (R19 in *Er*NaR) is highlighted blue. **C.** The RSB region of KR2 in the ground (left, yellow) and O (right, blue) states of the photocycle and the role of L74. Distance from L74 to the nearest water molecule in the Schiff base cavity is indicated with a red arrow (left) and is given in bold italic. The flipping motion of the L74 side chain upon sodium binding in the O state is also indicated with a red arrow (right). The sodium ion is shown with the purple sphere. **D.** Interprotomeric sodium binding site in KR2. Overall view of the KR2 pentamer from the extracellular side (left) and detailed view of the site (right). Two neighboring protomers are colored yellow and green. The BC loop is colored dark blue. The sodium ion (purple sphere) coordination is indicated with black dashed lines.

Most of the investigated NDQ rhodopsins belong to Subgroup 1. Our analysis shows that only one protein, a sodium pump from *Salinarimonas rosea* DSM21201 (*Sr*NaR), from Subgroup 2 was studied^6^. *Sr*NaR has been shown to differ from other NaRs. To date, the available data on *Sr*NaR and the members of Subgroup 2 are very limited. Therefore, to study the properties of the members of Subgroup 2, we selected another representative, an NDQ rhodopsin from *Erythrobacter sp.* HL-111 (*Er*NaR), and performed its thorough characterization. This allowed us to compare the properties and organization of *Er*NaR with known sodium pumps like KR2.

### Functional characterization of *Er*NaR

To study *Er*NaR in mammalian cells, we cloned a human-codon optimized gene of *Er*NaR to the previously published expression cassette that contained the enhanced yellow fluorescent protein (EYFP), membrane trafficking signal (TS) and endoplasmic reticulum (ES) export signal from potassium channel Kir2.1 and N-terminal part of channelrhodopsin (C2C1)^22^. We transiently expressed *Er*NaR in this construct in HEK293T cells and evaluated its localization by confocal imaging. Subcellular expression of *Er*NaR was predominantly confined to the plasma membrane, with low to none at all amounts of protein in intracellular compartments (Fig. 2A).

**Fig. 2.**
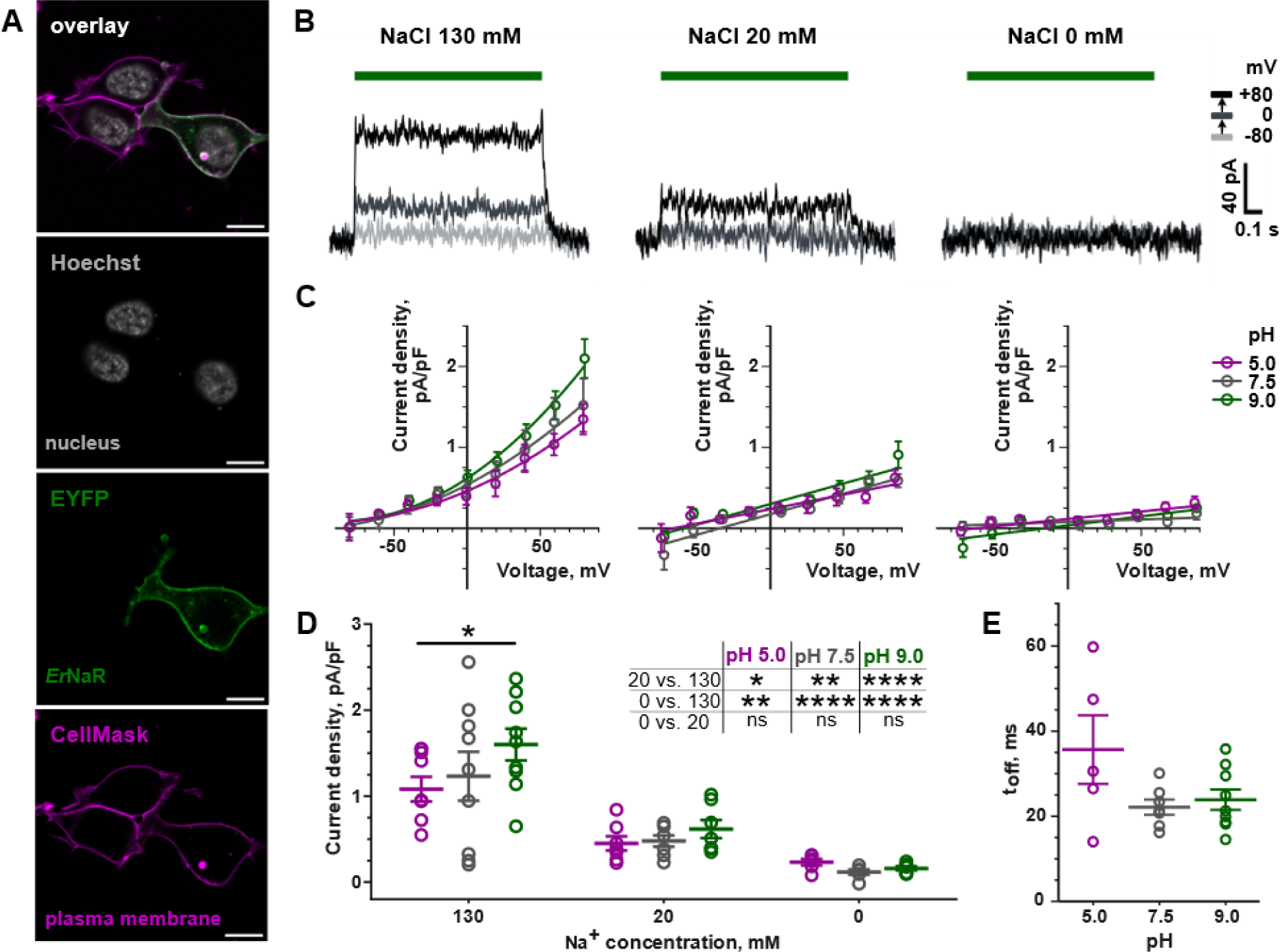
Functional characterization of *Er*NaR in HEK293T cells. **A.** Representative confocal images of HEK293T cell expressing *Er*NaR. Fluorescence of EYFP, fused to *Er*NaR, is shown in green; the plasma membrane stain CellMask - in magenta; nucleus stain Hoechst 33342 - in grey. The co-localization of EYFP and CellMask appears in white. Scale bars, 10 µm. **B.** Representative photocurrents of *Er*NaR recorded from HEK293T cells at 130 mM (left), 20 mM (middle) and 0 mM (right) intracellular [Na^+^]_i_ and pH_i_ 7.5. **C.** Voltage dependence of the stationary photocurrents of *Er*NaR at intracellular [Na^+^]_i_ 130 mM (left), 20 mM (middle) and 0 mM (right) and different pH_i_ values (LJP-corrected; normalized to respective cell capacitance; mean ± SEM of n = 6-9 cells). **D.** Stationary photocurrents of *Er*NaR at +60 mV, normalized to respective cell capacitance (mean ± SEM and individual data points of n = 6-9 cells). Data were extracted from the recordings at different intracellular [Na^+^]_i_ and pH_i_ described in **C.**. Normalized currents were analyzed using two-way ANOVA with two Turkey’s multiple comparisons tests – for the effect of pH_i_ at fixed [Na^+^]_i_ (depicted on the graph, ns is not shown) and for the effect of [Na^+^]_i_ at fixed pH_i_ (inset table). **E.** Kinetics of photocurrent decay upon light-off at 130 mM intracellular [Na^+^]_i_ and different pH_i_ (mean ± SEM and individual data points of n = 5-8 cells). The time constant (t_off_) was determined by monoexponential fit of photocurrent decay at holding voltage +80 mV. Data were analyzed using Kruskal-Wallis test with Dunn’s multiple comparisons test (all ns). **B-E.** All patch-clamp experiments were conducted at 110 mM extracellular [Na^+^]_e_, pH_e_ 7.5; LED light with maximum at 550 nm was applied for 1 s at 34.3 mW/mm² irradiance. ns – not significant (*P* > 0.05), \**P* < 0.05, \*\**P* < 0.01, \*\*\**P* < 0.001 and \*\*\*\**P* < 0.0001.

Next, we used the whole cell patch-clamp technique to study the functional properties of *Er*NaR. Expecting *Er*NaR to be an outward sodium or proton pump, we measured the photocurrents at the same extracellular pH (pH_e_) 7.5 and 110 mM [Na^+^]_e_ while varying intracellular pH (pH_i_) and [Na^+^]_i_. Indeed, *Er*NaR appeared to be an outwardly directed sodium pump, with photocurrent highly dependent on intracellular [Na^+^]_i_ (Fig. 2B-D). Notably, *Er*NaR showed a profound nonlinear voltage dependence at 130 mM [Na^+^]_i_, with near-zero currents in negative voltages (Fig. 2C, left). A voltage dependence was also reported in KR2^22^, but it was not as pronounced as observed in *Er*NaR.

To assess the capability of *Er*NaR to pump H^+^, for each [Na^+^]_i_ we tested three pH_i_ values (5.0, 7.5, and 9.0) (Fig. 2C). In the presence of sodium in the intracellular solution (20 mM and 130 mM [Na^+^]_i_) we observed photocurrents at all studied pH_i_. A change in pH_i_ from acidic (5.0) to alkaline (9.0) slightly increased the amplitude of photocurrent (Fig. 2C, left and middle plots). While at 20 mM [Na^+^]_i_ this effect was not significant, at 130 mM [Na^+^]_i_ the difference reached statistical significance (*P*=0.0349) when the photocurrents were compared at +60 mV (Fig. 2D). In addition, at 130 mM [Na^+^]_i_ we determined the characteristic time of the photocurrent decay after the light was switched off (t_off,_ monoexponential fit) in all tested pH_i_ values (Fig. 2E). Acidic intracellular conditions led to slightly decelerated off-kinetics of *Er*NaR (statistically not significant).

In the absence of intracellular sodium, *Er*NaR exhibited immeasurably low photocurrents in all tested pH_i_. Under similar experimental conditions, when sodium was removed from the intracellular solution, KR2 was shown to pump protons^22^. In *Er*NaR, the proton currents were negligible even at pH_i_ 5.0, suggesting that *Er*NaR and likely other members of Subgroup 2 of the NDQ rhodopsins are more selective to sodium than representatives of Subgroup 1. Besides, the difference in photocurrent amplitude at +60 mV was statistically significant between 130 mM and 20 mM [Na^+^]_i_ (*P=*0.0142*, P=*0.0028 and *P<*0.0001 at pH_i_ 5.0, 7.5 and 9.0, respectively) and between 130 mM and 0 mM [Na^+^]_i_ (*P=*0.0012*, P<*0.0001 and *P<*0.0001 at pH_i_ 5.0, 7.5 and 9.0, respectively) (Fig. 2D). However, the difference between 20 mM and 0 mM [Na^+^]_i_ was not significant in all pH_i_, suggesting that *Er*NaR might require higher intracellular sodium concentrations to successfully function.

### Spectroscopy of *Er*NaR

Next, we studied the spectroscopic properties of detergent-solubilized *Er*NaR. In contrast to KR2^1^ and many other microbial rhodopsins^1,23^, *Er*NaR undergoes only a small spectral red-shift of 3 nm upon acidification from pH 8.0 (λ_max_ 535.5 nm) to pH 4.3 (λ_max_ 538.5 nm) (Fig. 3A). An additional 6.5 nm red-shift is observed upon further acidification to pH 2.3 (λ_max_ 545.0 nm) (Fig. S1). Usually, pH dependent spectral changes of the dark state absorption spectrum are caused by altered protonation states of either the RSB or its counterion^1,24,25^. For KR2, it was shown that the main counterion is the aspartate (D116) of the characteristic NDQ motif, which gets protonated upon acidification and the absorption spectra undergoes notable red-shift by more than 20 nm already at pH 4.3^1^. The protonation of D116 leads to the altered photocycle kinetics and strong decrease in ion-pumping activity^1,12^. In *Er*NaR, D105 residue is located at this position, which is also the main RSB counterion directly H-bonded to the Schiff base in the resting state. Based on the spectroscopy data showing almost no spectral shift upon acidification, we conclude that D105 in *Er*NaR remains deprotonated in a wide range of pH values. This could be explained by the presence of an additional glutamate E64 in the RSB region of this rhodopsin. The pH titration of the dark state of the *Er*NaR absorption spectrum (Fig. S1) yielded two pK_a_ values (pK_a,1_ = 3.34 ± 0.04 (Fig. S1B) and pK_a,2_ = 5.71 ± 0.19 (Fig. S1C)). It should be noted that in the case of detergent-solubilized KR2, the protein dissociates to monomers upon acidification to pH 4.3^12^, which is also most likely reflected in the spectral properties of the rhodopsin^1^. On the contrary, the detergent-solubilized *Er*NaR remains pentameric at both pH 4.3 and 8.0 as clearly observed in size-exclusion chromatography and cryo-EM experiments. Thus, the observed minor spectral shift is the feature of the functionally-relevant oligomeric form of *Er*NaR.

**Fig. 3.**
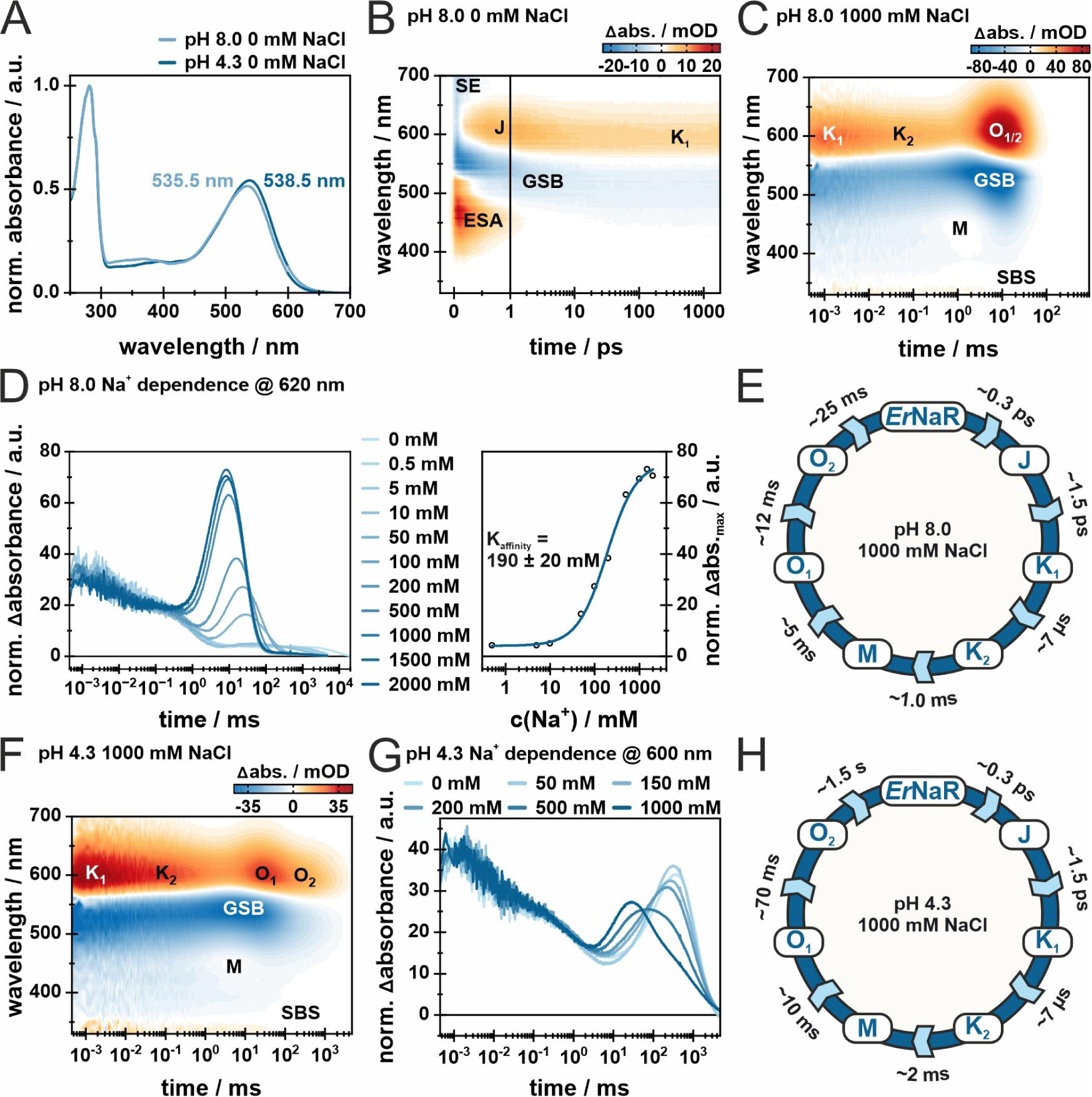
Spectroscopic characterization of *Er*NaR. **A.** Normalized absorption spectra of *Er*NaR at acidic (pH 4.3) and neutral (pH 8.0) conditions. The spectra have been normalized to the absorbance at 280 nm. **B.** 2D-contour plot of a fs-TA measurement of *Er*NaR (pH 8.0, 0 mM NaCl). The timescale is linear until 1 ps and logarithmic afterwards. The signal amplitude is color coded as follows: positive (red), no (white) and negative (blue) abs. The mentioned abbreviations represent excited-state absorption (ESA), ground-state bleach (GSB) and stimulated emission (SE). **C.** 2D-contour plot of a flash photolysis measurement of *Er*NaR (pH 8.0, 1000 mM NaCl). **D.** Sodium dependence of the transient at 620 nm, indicative of the O_1/2_-intermediate, at pH 8.0. Transients have been normalized to have the same Δabs. at 0.1 ms. The absorption maximum of the O_1/2_-intermediate was plotted against the sodium concentration and the resulting data was fitted with the Hill equation to obtain the sodium dependence of the O_1/2_-intermediate. **E.** Schematic model of the *Er*NaR photocycle at pH 8.0, 1000 mM NaCl. **F.** 2D-contour plot of a flash photolysis measurement of *Er*NaR (pH 4.3, 1000 mM NaCl). **G.** Sodium dependence of the transient at 600 nm, representative for the O_1/2_-intermediates, at pH 4.3. Transients have been normalized to have the same Dabs. at 0.1 ms. **H.** Schematic model of the *Er*NaR photocycle at pH 4.3, 1000 mM NaCl.

A similar weak pH dependence was found for the ultrafast dynamics of *Er*NaR. For all measured conditions (pH 8.0 and 4.3), the ultrafast pattern looks similar to what was already shown for other microbial rhodopsins at neutral to alkaline pH values (Fig.3B and Fig. S2)^23,26–29^. Namely, after photo-excitation, the excited state decays on a sub-ps timescale and the formation of a hot ground state intermediate at ∼610 nm is observed. This intermediate is commonly termed as J and relaxes to form the first stable red-shifted photoproduct K_1_ at ∼590 nm described by the lifetime distribution in the 1-4 ps range (Fig. S2A).

We also studied the photocycle kinetics of *Er*NaR on the ns to s timescale under various conditions using flash photolysis. At pH 8.0, the photocycle, as well as the impact of different sodium concentrations on the *Er*NaR, are similar to what was observed for KR2^1,30,31^. The K_1_-intermediate at the end of the ultrafast measurement (1.8 ns), remains present at the beginning of the flash photolysis time scale (450 ns), showing that K_1_ is populated throughout the ns timescale and no photocycle intermediate is missed due to the experimental time gap (Fig. 3B,C). K_1_ decays within the early µs-range (lifetime distribution in the range of 5-10 µs) to form the K_2_-intermediate. This intermediate state is considered to be K-like because of the more pronounced red-shift compared to the known L-intermediates of other microbial rhodopsins^30,31^. The transition to the blue-shifted M-intermediate then occurs within ∼1 ms. In the absence of sodium, M is the last observed intermediate of the photocycle and the protein subsequently relaxes back to the parent state, represented in the lifetime distributions centered at 5-7 s (Fig. S3A). In the presence of a sufficient amount of sodium ions, however, the duration of M is significantly shortened and, as was observed also for KR2^1^, a strong signal assigned to the red-shifted O-intermediate rises (Fig. 3D (left)) within 2-10 ms (Fig. S3C). Comparison of the maximum signal intensity of the O-states for different NaCl concentrations allowed the determination of K_affinity_ = 190 ± 20 mM for *Er*NaR, demonstrating that high sodium concentrations are required to observe the “sodium-pumping mode” (Fig. 3D). At 100 mM NaCl, *Er*NaR shows an equilibrium of M- and O-intermediates (Fig. S3B), indicative for mixed kinetics of one subpopulation likely undergoing the “sodium-pumping mode” photocycle while the other subpopulation undergoes a long-living M-intermediate before directly relaxing back to the parent state. In the case of the sodium-pumping regime, an additional lifetime (10-25 ms) is needed to fully describe the O-intermediate kinetics indicated by the spectral shift within the positive lifetime distribution in the range of 10-55 ms (Fig. S3C). As it was mentioned earlier, the existence of one or two O-intermediate components and the respective interpretation is currently under debate^14,15,17^. *Er*NaR shows two distinct O-intermediate components (O_1_ and O_2_), with O_2_ blue-shifted compared to O_1_ and - according to the proposed retinal configuration marker band at ∼335 nm^32^ - O_1_ contains 13-*cis* retinal, while O_2_ has all-*trans* retinal (Fig. S3). These observed experimental results are in line with the findings and interpretation of Fujisawa et al.^17^ for *Ia*NaR.

At pH 4.3, the photocycle kinetics of *Er*NaR differs from that at pH 8.0. The kinetics of the early photocycle intermediates up to the formation of M under all tested conditions are similar to that observed at pH 8.0. The M-intermediate and the two O-intermediates again exhibit sodium dependence but at pH 4.3 the effects are clearly weaker. For instance, at pH 4.3 the M state under all conditions shows a weak absorbance and a fast decay, similar to that found for *Er*NaR at pH 8.0 at the sodium concentration of 1000 mM. Such a fast decay of the M state, which corresponds to the reprotonation of the RSB, can be explained by the higher proton concentration at acidic conditions and was also observed for KR2^33^.

While for KR2, the distinct red-shifted O-intermediate is present only in the sodium-pumping mode and was directly associated with the sodium binding during the photocycle^1,17^, this must not necessarily be the case for *Er*NaR. Indeed, the O_1_ and O_2_ states were observed for *Er*NaR at pH 4.3 for every sodium concentration including the sodium-free conditions. The spectral shapes of the O_1_ and O_2_ intermediates differ from those at pH 8.0. At pH 4.3, the spectra of both O states are clearly blue-shifted compared to those at pH 8.0, while the red-shift was more expected (Fig. S4,S5)^24,30,32,34^. This indicates that in the late stages of the *Er*NaR photocycle the environment in the close vicinity to the retinal chromophore is likely significantly different at pH 4.3 and 8.0.

With increasing sodium concentration at pH 4.3, the kinetics of the two O-intermediates are influenced differently, leading to a spectral and temporal separation of O_1_ and O_2_ at 1000 mM NaCl (Fig. 1G). Additionally, the retinal isomerization marker band in the near-UV (SBS)^32^ indicates that the retinal conformation is different in the two O-intermediates, namely a 13-*cis* configuration is suggested for O_1_ and all-*trans* configuration for O_2_, respectively (Fig. 3). To exclude possible ionic strength effects, measurements with 1000 mM KCl and 1000 mM N-Methyl-D-glucamine (NMG), an organic monovalent cation that is commonly used to replace sodium ions in electrophysiological experiments^35^, were performed. The measurements at pH 4.3 with 1000 mM KCl showed a photocycle very similar to the one obtained for 0 mM and 100 mM NaCl at pH 4.3 (compare Fig. S4A,B and Fig. S6B) lacking the clear spectral differences observed for 1000 mM NaCl (Fig. S4C and S6B). With 1000 mM NMG at pH 4.3, the photocycle is very similar to the one with 1000 mM KCl and 0 mM NaCl at pH 4.3. The obtained lifetime densities for 1000 mM KCl and 1000 mM NMG are in good agreement as well (compare Fig. S4A and S6). Therefore, this allowed us to rationalize that under acidic conditions the observed separation of the O_1_ and O_2_ states in response to the increase of sodium concentration is a direct result of the sodium binding close to the RSB.

### Cryo-EM structure of the pentameric *Er*NaR

In order to investigate the molecular basis of the unique functional and spectral properties of *Er*NaR and the effect of Leu to Glu replacement in rhodopsins, we used single-particle cryo-electron microscopy (cryo-EM) to determine the structure of the protein in different conditions. We obtained the structures of *Er*NaR at acidic (4.3) and neutral (8.0) pH values at the resolution of 2.50 and 2.63 Å, respectively (Fig. S7). Cryo-EM maps allowed us to model residues 1-272 of *Er*NaR. The overall organization of the pentamer is similar to that of KR2 and other MRs (Fig. 4A; Fig. S7). Namely, the nearby protomers interact via helices A and B and BC loop containing a β-sheet (Fig. S8).

**Fig. 4.**
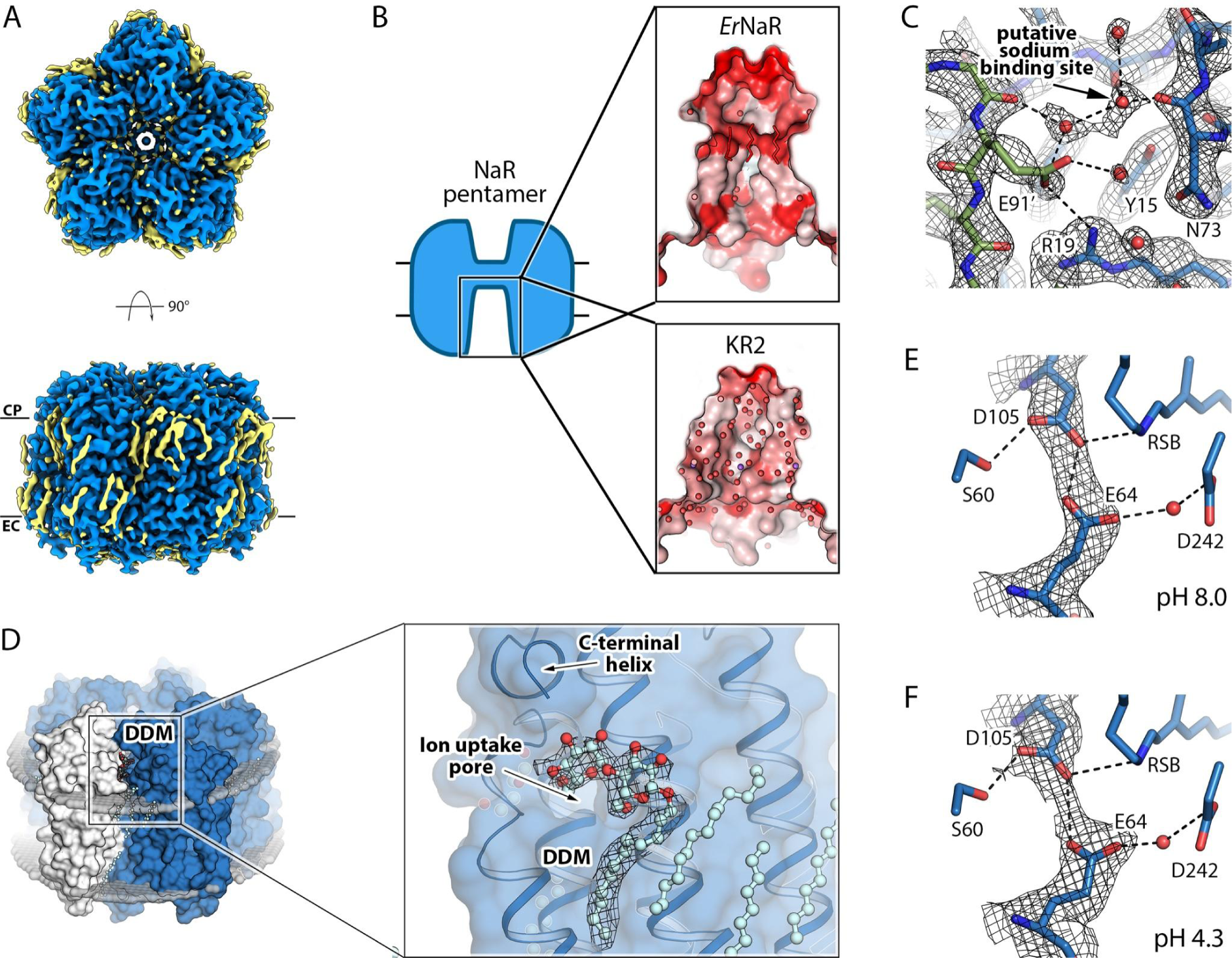
Cryo-EM structures of *Er*NaR. **A.** Overall view of the *Er*NaR pentamer from the extracellular (top) and along the membrane plane (bottom). **B.** Comparison of the hydrophobicity of the concave aqueous basic in the center of the *Er*NaR and KR2 pentamers. **C.** The region of interprotomeric sodium binding site (according to the KR2 structure) absent in *Er*NaR. The cryo-EM map (pH 8.0) is shown with black mesh. **D.** The DDM molecule in the cryo-EM structures of *Er*NaR. 5 molecules of DDM are located between neighboring protomers at the cytoplasmic side of the protein. The polar head of DDM is located next to the entrance to the ion uptake cavity and interacts with the C-terminal helix found in *Er*NaR. **E.** The central region of the *Er*NaR protomer at pH 8.0. The cryo-EM map is shown with black mesh and indicates direct interaction between D105 and E64. **F.** The central region of the *Er*NaR protomer at pH 4.3. The cryo-EM map is shown with black mesh and indicates direct interaction between D105 and E64.

At the same time, the concavity in the central part of the pentamer at the extracellular side is organized differently in *Er*NaR and KR2 (Fig. 4B). In general, while in KR2 this region is polar and interacts with numerous water molecules identified using X-ray crystallography (PDB ID: 6YC3^19^), in *Er*NaR it is more hydrophobic (Fig. 4B). Moreover, the cryo-EM maps reveal the presence of fragments of hydrophobic chains in this region of *Er*NaR (Fig. 4B).

Notably, the cryo-EM data strongly suggest that the interprotomeric sodium binding site found in the KR2 pentamer^9^ is absent in *Er*NaR (Fig. 4C). Although the maps show the spherical density at a position similar to that of the sodium ion in KR2 (Fig. 1D, 4C), the coordination is unlikely to favor sodium binding. In KR2 the sodium binding site is formed by two neighboring protomers; namely, by the side chains of D102’ and Y25 as well as carbonyl oxygens of T83 and F86 and two water molecules (Fig. 1D). In *Er*NaR, E91 (corresponding to D102 in KR2) is located 2.5 Å further from the position of the probable sodium binding site and thus is unlikely to coordinate the ion (Fig. 4C). Moreover, E91 directly interacts with the positively charged R19, located within only 5.5 Å from the possible sodium binding site (Fig. 4C). The end of BC loop of *Er*NaR is also arranged slightly differently from that of KR2 (Fig. S8). The sequence alignment also indicates that the region of the interprotomeric sodium binding site is organized differently in Subgroups 1 and 2 of NDQ rhodopsins (Fig. 1B, right). Thus, we speculate that the interprotomeric sodium binding site found in KR2 is likely a feature of only Subgroup 1 but not Subgroup 2 of NDQ rhodopsins.

Another feature of *Er*NaR is the presence of a detergent (n-Dodecyl-beta-Maltoside, DDM) molecule in the cleft between the rhodopsin protomers (Fig. 4D). The DDM molecule is found at the cytoplasmic leaflet of the membrane and its polar head is located near the pore in the *Er*NaR surface likely serving as the entrance for sodium ions (Fig. 4D). The polar head also interacts with the C-terminus of the protein (Fig. 4D). Nevertheless, the DDM molecule does not block the pore.

Although the quality of the cryo-EM map is sufficient to place the side chains of amino acid residues as well as protein-associated water molecules, it is still limited and does not allow us to precisely determine the distances between the functional groups of rhodopsin.This hampers the understanding of the molecular mechanisms of *Er*NaR. For instance, while the cryo-EM data clearly show a direct interaction between the D105 residue of the characteristic NDQ motif with the newly identified E64 residue of *Er*NaR at both pH 4.3 and 8.0, the map regions corresponding to these residues merge together and individual positions cannot be resolved (Fig. 4E,F). Nevertheless, this might be indicative that there is a rather short H-bond between these two residues.

### The internal organization of the *Er*NaR protomer

In order to resolve details of the internal organization of *Er*NaR, we crystallized the protein using *in meso* approach and determined its structures at pH 4.6 and 8.8 at 1.7 Å resolution using X-ray crystallography. The crystals originally appeared at pH 4.6 and contained a monomer of *Er*NaR in the asymmetric unit. To obtain high-resolution structure of *Er*NaR at high pH, we soaked the crystals in the buffer solution with pH 8.8 (see Methods for details). The structures of the rhodopsin at pH 4.6 and 8.8 appeared nearly identical (RMSD of 0.06 Å). The structure of the *Er*NaR protomer obtained using X-ray crystallography is also very similar to that in the pentameric cryo-EM structure (RMSD of overall structures of 0.5 Å). Since the crystal structure is similar to that obtained with cryo-EM but reveals more details on the internal organization of *Er*NaR, we used it for the further analysis.

The overall protomer organization of *Er*NaR is similar to that of KR2 (Fig. 5A; Fig. S8C,D) with a large cavity at the cytoplasmic part of both *Er*NaR and KR2, likely acting as an ion uptake cavity (Fig. 5A,B). In the central region, D105 (D116 in KR2) is directly H-bonded to the RSB (Fig. 5C). The retinal cofactor is in the all-*trans* configuration in the resting state of *Er*NaR (Fig. 5A). The internal extracellular region of *Er*NaR is separated from the RSB region with R98 (analog of R109 in KR2) and comprises numerous polar residues including the E5-E149-R234 triad, organized almost identically to the E11-E160-R243 triad of KR2 (Fig. 5A).

**Fig. 5.**
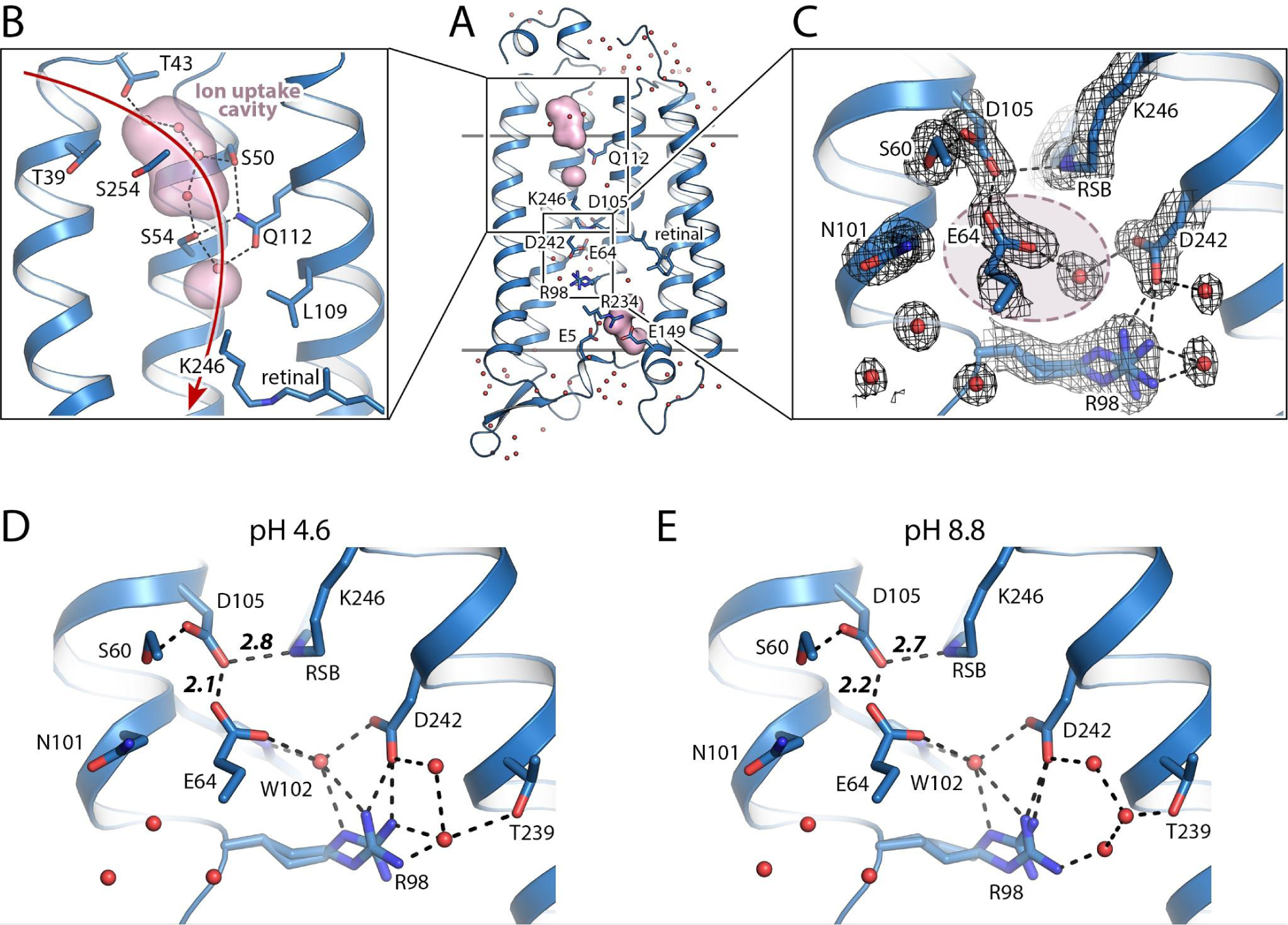
Crystal structures of *Er*NaR. **A.** Overall side view of the *Er*NaR monomer. Hydrophobic/hydrophilic membrane core boundaries are shown with gray lines. **B.** Detailed view of the cytoplasmic internal part of *Er*NaR. Cavities were calculated using HOLLOW and are shown with a pink surface. Red arrow indicates putative sodium uptake pathway. **C.** The RSB region of *Er*NaR at pH 4.6. 2Fo-Fc electron density maps are contoured at the level of 1.0σ and are shown with black mesh. H-bonds are shown with black dashed lines. Light-red area indicates the region of the Schiff base cavity found in KR2 and absent in *Er*NaR. **D.** The detailed view of the RSB region of *Er*NaR at pH 4.6. **E.** The detailed view of the RSB region of *Er*NaR at pH 8.8. H-bonds are shown with black dashed lines. The lengths of the key H-bonds between the RSB and D105 and D105 and E64 are given in Å and shown with bold italic numbers.

Despite some similarities, the protomer structure of *Er*NaR differs from that of KR2 in several aspects. First, the N-terminus is shorter in *Er*NaR and the N-terminal α-helix has only one turn compared to two turns in KR2 (Fig. S8C). Nevertheless, the N-terminus is capping the inside of *Er*NaR (Fig. 5A, S8C). Also, an additional small C-terminal α-helix was found in *Er*NaR, comprising residues 267-272 of the protein (Fig. S8D).

Second, the cytoplasmic ends of helices D, E, and F are shifted by more than 3 Å in *Er*NaR compared to those of KR2 (Fig. S8D). This might be related to the notable differences in the length and organization of the CD and EF loops of the two rhodopsins (Fig. S8D). However, these shifts of the helices are not reflected in the internal organization of the cytoplasmic part of *Er*NaR. In particular, the ion uptake cavities of *Er*NaR and KR2 are highly similar.

Some differences between KR2 and *Er*NaR were also found in the extracellular side of the proteins. For example, there is a leucine (L68) in *Er*NaR at the position of Q78 in KR2, which was shown to be critical for sodium release from the latter rhodopsin^19^. However, the Q78L mutation of KR2 has no effect on sodium pumping; therefore, we suggest that this natural substitution found in *Er*NaR is in line with the earlier suggested mechanism of sodium release from the NDQ rhodopsins^19^.

Another noteworthy difference between *Er*NaR and KR2 was found in the RSB region. E64, a unique characteristic residue of *Er*NaR and Subgroup 2 of NDQ rhodopsins, is pointed towards the RSB and interacts directly with D105 (D116 in KR2) (Fig. 5C). As a result, a large Schiff base cavity found in the ground state of KR2 in the pentameric form is absent in *Er*NaR and only one water molecule is found in this region (Fig. 5A,C). It is coordinated by E64, R98, W102, and D242 (Fig. 5C,D). At the same time, the N101 side chain (N112 in KR2) is pointed towards the oligomerization interface similar to that found in the ground state of pentameric KR2^12^ (Fig. 5C,D).

The interaction between D105 and E64 has not been observed before and is an intriguing feature of *Er*NaR. Significantly higher resolution of the X-ray crystallography data (1.7 Å vs 2.5 Å for the cryo-EM data) allowed us to accurately position the side chains of E64 and D105 at both pH 4.6 and 8.8 (Fig. 5C). The distance between the nearest oxygens of E64 and D105 was only 2.1 and 2.2 Å at pH 4.6 and 8.8, respectively, which was extremely short for a donor-acceptor distance in a normal H-bond and strongly suggests a low-barrier H-bond (LBHB) between the residues (Fig. 5D,E). The E64 side chain was also connected to the second aspartic acid counterion of the RSB, D242, through a water-mediated H-bond chain (Fig. 5C-E).

In order to get insights into the protonation states of carboxylic residues in close vicinity to the RSB and to gain more information on the strong interaction between E64 and D105 we performed hybrid quantum mechanics/molecular mechanics (QM/MM) simulations using the crystal structure of *Er*NaR. In particular, the protonation state of the E64 residue is of high interest as this residue is only found in Subgroup 2 of NDQ rhodopsins. Thus, we probed various protonation states of the counterion complex. For each of the three counterions we have considered one of four proton orientations as shown on Fig. S9. In total 61 protonation patterns were considered (1 with zero-protonation, 12 single-protonation and 48 double-protonation). For all protonation pattern structure optimization was followed up by the calculation of vertical excitation energies (Fig. S9A).

We found that the spectral and structural properties of *Er*NaR can be explained by system 6 with two protonated counterions, namely, E64 and D242 (Fig. S9A,B). This model also shows the lowest ground state energy with respect to all considered protonation models. The models with one protonated counterion appear significantly blue-shifted compared to those with two protonated counterions. The protonation of D105 would cause the red shift of the spectra by more than 60 nm, but its ground state energy is too high. Moreover, it would break the D105-RSB salt bridge, which is clearly observed in the high-resolution structures. Thus, we conclude that in agreement with the spectroscopic and structural data on *Er*NaR, D105 remains deprotonated in a wide range of pH values, while E64 is likely protonated even at a pH as high as 8.8. In the QM/MM-refined structure (Fig. S9D) the distance between D105 and E64 is 2.5 Å. In order to analyze the nature of the interaction we performed the energy decomposition analysis using Zero-Order Symmetry Adapted Perturbation Theory (SAPT0). The SAPT0 analysis shows a significant repulsion, but it was overcompensated by electrostatics, polarization, and dispersion interactions (Fig. S9B).

As it was proposed from the spectroscopy data, the newly identified E64 residue in the helix B of *Er*NaR likely maintains low pKa of the main RSB counterion D105. Structural data and QM/MM simulations suggest the following mechanism of the E64 influence on the pKa of D105 with at least two components: (1) an unusually short H-bond to the protonated E64 affecting the electrochemical properties of D105; (2) the tight integration of the E64 side chain into the H-bond network of the RSB region additionally stabilizes the overall conformation associated with the deprotonated form of D105. The LBHB between E64 and D105 means that the E64-D105 pair might also share a proton in the ground state of *Er*NaR in a similar way that shown for the proton release group of BR^36^.

### Comparison of the two subgroups of NaRs

The bioinformatic analysis of the clade of NDQ rhodopsins revealed two major subgroups. Our functional, spectroscopical, and structural data on *Er*NaR from Subgroup 2 presented here and the previously reported data on KR2 as well as other NDQ rhodopsins of Subgroup 1 allowed us to compare the properties and molecular mechanisms of these two subgroups of NaRs.

Electrophysiology demonstrated that *Er*NaR is capable of active transport of sodium across the membrane in a wide range of pH values, including acidic pH, which is in contrast to the functional properties of KR2. Indeed, in KR2, as well as in other members of Subgroup 1, the pH decrease leads to the protonation of the main RSB counterion, aspartic acid of the characteristic NDQ motif^1,3^. This protonation results in the altered photocycle and a dramatic decrease in ion-pumping activity^1,12^. On the contrary, the members of Subgroup 2 lack such pH dependence both on spectra and sodium-pumping efficiency. In addition, proton pumping by *Er*NaR was negligible at all tested pH values and sodium concentrations. This is also in contrast to KR2^1^. Therefore, we suggest that the presence of an additional carboxylic residue in close proximity to the RSB in the rhodopsins of Subgroup 2 might lead to their higher selectivity to sodium ions over protons and the possible absence of the proton-pumping mode.

Our kinetics studies demonstrated both common features and fundamental differences between Subgroup 1 and Subgroup 2 of NDQ rhodopsins. Clearly, the acceleration of the M state decay upon the increase of either sodium or proton concentration shows that both types of ions can be uptaken by *Er*NaR in a competitive manner, similar to what was found for KR2^33^. However, since the proton-pumping activity of *Er*NaR is weak compared to KR2, the further proton transition steps likely proceed differently in Subgroups 1 and 2. Namely, we suggest that in Subgroup 1 the proton is released to the extracellular space at the end of the photocycle, while in *Er*NaR it is likely released back towards the cytoplasm through the ion uptake cavity.

While the conformation of the RSB region in the resting state is different in *Er*NaR and KR2, there is a similarity in the orientation of N101 and N112 of the characteristic NDQ motif in *Er*NaR and KR2, respectively (Fig. 6). The asparagine is oriented outside of the protein protomer towards the pentameric interface in the resting state of both rhodopsins. For KR2, this conformation was directly demonstrated in 2019 but was originally proposed back in 2016 and was named “expanded” as it is characterized by a large water-filled cavity near the RSB^12,37^. The expanded conformation was only found at neutral pH values, when the rhodopsin works as a sodium pump^12^. Thus, this conformation was assigned to the functional form of the protein. In *Er*NaR, the conformation of the RSB region lacks the large Schiff base cavity but with respect to the N101 orientation is similar to the expanded one of KR2. This allows us to conclude that in the resting state of all NDQ rhodopsins under sodium-pumping conditions, including both Subgroups 1 and 2, the common characteristic feature is the orientation of asparagine of the functional motif outside of the protomer towards the oligomerization interface. We named this common conformation as the “N-out” one (Fig. 6). Consequently, the conformation of NDQ rhodopsin with the asparagine oriented inside the protomer should be named “N-in” (Fig. 6).

**Fig. 6.**
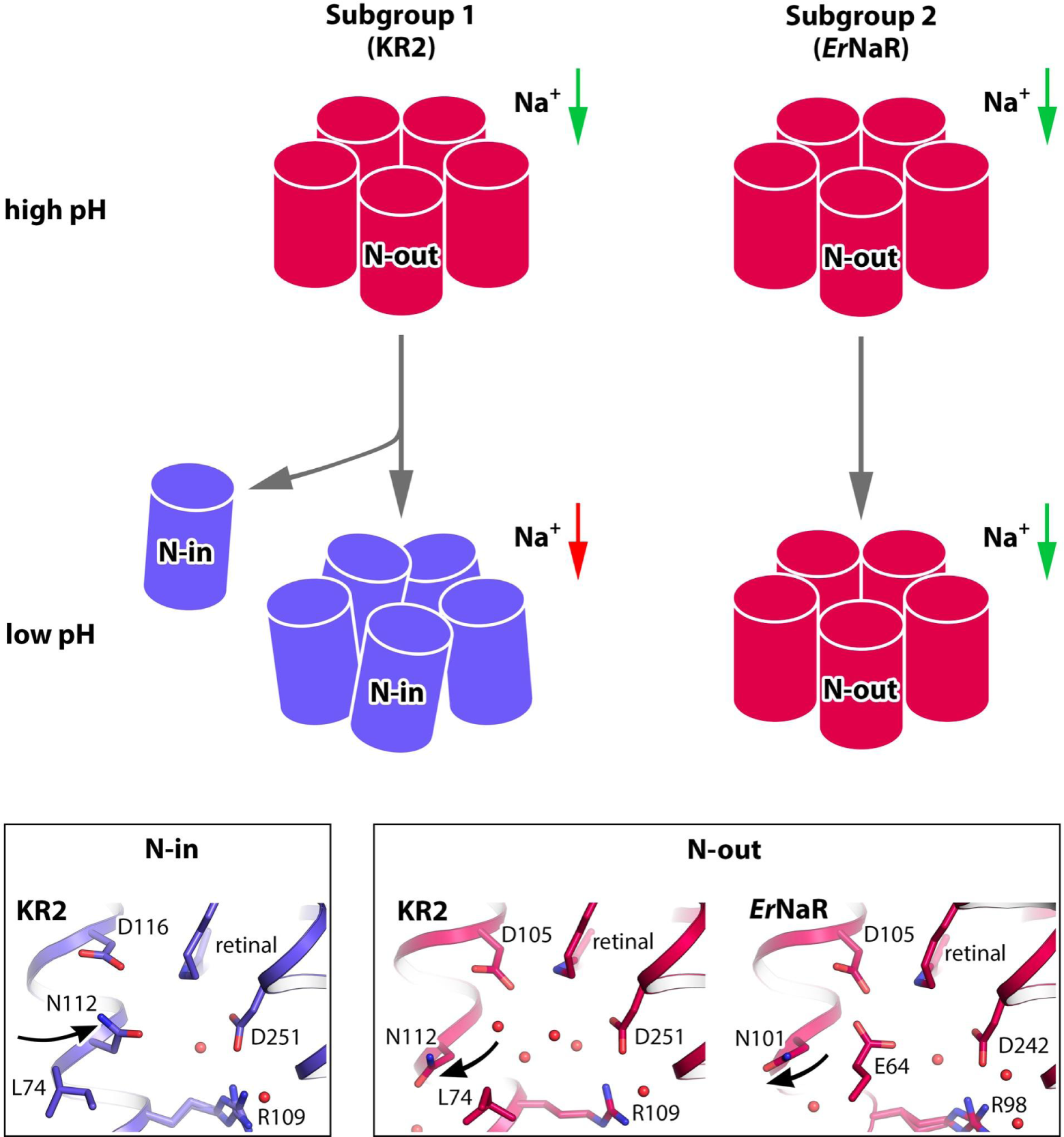
**N**-in and N-out conformations of the NDQ rhodopsins. The scheme of the difference in pH dependence of the spectral properties (spectral shift upon acidification in Subgroup 1 is shown with the change of the color from pink to blue), the pentamer stability (the disturbed pentameric assembly at low pH is shown for KR2 schematically by disoriented protomers; partial dissociation of the pentamers into protomers is also indicated by an additional gray arrow), and the conformation between Subgroup 1 and 2 of NDQ rhodopsins (N-in and N-out conformations of the active centers). Green arrows indicate sodium transport across the membrane by NaRs. Red arrow indicates the absence of sodium transport. The detailed view of the RSB regions of KR2 and *Er*NaR in N-in and N-out conformations is given in the bottom panels. Black arrows indicate orientation of Asn residue of the NDQ motif different in N-in and N-out conformations.

As shown for KR2, the N-out conformation is provided by the pentameric assembly^12^. The N112 side chain is H-bonded to the residues of the nearby rhodopsin molecule when flipped outside of the protomer. At low pH values the pentameric assembly of KR2 is disturbed and oligomers partially dissociate into monomers in detergent micelles^12^ (Fig. 6). In pentamers of KR2 at low pH only the N-in conformation was found^12^. Furthermore, in the monomer, the N-in conformation was always observed and the major reason for that was suggested to be the absence of the N112 stabilization^12^. In contrast to KR2, the pentameric assembly of *Er*NaR is more stable and pH independent. *Er*NaR also adapts the N-out conformation at both high and low pH values (Fig. 6). Even in the monomeric form of *Er*NaR formed exclusively in crystals the N-out conformation is present at both acidic and alkaline pH. In summary, we suggest that the conformations of the residues comprising the characteristic NDQ motif (N101, D105, Q112 in *Er*NaR and N112, D116, Q123 in KR2) are similar in the ground states of both proteins, which is likely a common feature of the members of both Subgroups 1 and 2. At the same time, this N-out conformation seems to be much more stable in the proteins of Subgroup 2 than of Subgroup 1, and is independent of pH and oligomeric state.

Although the pentameric assembly is a common feature of all NDQ rhodopsins, the central areas of the pentamers of the members of Subgroups 1 and 2 are organized differently. The interprotomeric sodium binding site found in KR2 and thought to be present in all proteins belonging to Subgroup 1, is likely absent in *Er*NaR and the rest of Subgroup 2. The environment of the concave aqueous basin in the middle part of the *Er*NaR oligomer is more hydrophobic than that of KR2. The sodium release through this central region and a possible relay mechanism involving the surface-bound interprotomeric sodium ion in KR2 were proposed to lower the energetic barriers for the ion translocation against strong electrochemical gradients^19^. Thus, we suggest that sodium-release pathways and mechanisms might be different in Subgroups 1 and 2 of the NDQ rhodopsins.

## Conclusion

Constantly expanding gene databases allowed us to discover the naturally occurring variations within the NDQ rhodopsin clade of light-driven sodium pumps. Our findings demonstrated a new natural way of fine-tuning the active sodium transporters by the introduction of an additional carboxylic residue in close proximity to the retinal Schiff base. It results in (1) efficient sodium pumping at a wide range of pH values, (2) spectral stability, and (3) high ion selectivity, which could make the members of the newly highlighted subgroup of NDQ rhodopsins more useful for biotechnological applications. While the mechanistic insights on the resting state reported here shed more light on the NaRs organization, further structural investigations of their intermediate states are required to understand their molecular mechanisms. According to the reported results, there might be multiple mechanisms of light-driven sodium pumping by NDQ rhodopsins. Our study may be the basis for the upcoming time-resolved crystallography and/or cryo-EM as well as cryotrapping studies of the newly identified NDQ rhodopsins.

## METHODS

### Search for and phylogenetic analysis of the NDQ rhodopsins

To retrieve the NDQ rhodopsin genes from available sequence databases, jackhmmer search (N iterations = 5) was performed using the sequence of KR2 rhodopsin (UniProt ID: N0DKS8) as a template against the UniProtKB (2023-09-15), UniParc (2023-09-15), Genbank (2023-09-15), and MGnify (2023-09-15) databases. The 6-letter motifs for each protein of the output multiple sequence alignment (MSA) file, corresponding to the amino acid residues at the positions of R109, N112, D116, Q123, D251, and K216 of KR2 were retrieved using a custom Python script using jupyter notebook. The sequences possessing the RNDQDK motif were selected, filtered to contain more than 250 amino acid residues, and re-aligned separately using MUSCLE^38^. Duplicated sequences were removed using a custom Python script. The selected sequences were also manually inspected to contain full seven transmembrane α-helices. The maximum likelihood phylogenetic tree was built using IQ-TREE webserver^39^ (Number of bootstrap alignments: 1000) and visualized using iTOL (v6.8)^40^. For the building of the phylogenetic tree all sequences with identity higher than 90% were removed using CD-HIT software^67^. The full MSA file, 90% identity-filtered MSA file, and phylogenetic tree file are provided as Supplementary files. Two subgroups of NDQ rhodopsins were separated by using the presence/absence of the glutamate of the position of L74 in KR2 as a criterion.

### Cloning

*Er*NaR coding DNA was optimized for *E.coli* or human codons using GeneArt (Thermo Fisher Scientific). Genes were synthesized commercially (Eurofins). For protein expression and purification, the pET15b plasmid with 6xHis-tag at the C-terminal was used. For electrophysiological recordings, a human codon optimized *Er*NaR gene was cloned into the pcDNA3.1(-) vector between *Bam*HI and *Hind*III sites together with an N-terminal part of channelrhodopsin (C2C1), membrane trafficking signal (TS) and ER export signal (ES) from potassium channel Kir2.1 and enhanced yellow fluorescent protein (EYFP). C2C1 and TS-EYFP-ES were amplified from pEYFP-N1-eKR2^22^, which was a gift from Peter Hegemann (Addgene plasmid #115337). The final construct (C2C1-*Er*NaR-TS-EYFP-ES) was verified by sequencing. The plasmid is available from Addgene (#209851).

### HEK293T cell culture and transfection

The HEK293T cells (Thermo Fisher Scientific) were cultured at 37 °C with 5% CO_2_ in high-glucose Dulbecco’s Modified Eagle Medium (DMEM), containing 1 mM sodium pyruvate and 1X GlutaMAX, supplemented with 10% heat-inactivated fetal bovine serum, 100 U/ml penicillin and 100 µg/ml streptomycin (all Thermo Fisher Scientific). The cells were regularly tested and were free from mycoplasma. For electrophysiological recordings and confocal imaging cells were seeded on poly-l-ornithine-coated 24-well plates at a concentration of 1.5 × 10^5^ cells per well. After 24h, the cells were transiently transfected using 1.5 µg of Polyethylenimine MAX (PEI) (Polysciences) and 1 µg of plasmid per well in Opti-MEM (Thermo Fisher Scientific), supplemented with 1 µM of all-*trans*-retinal (Sigma). The transfection medium was replaced after 4–5 h with full culture medium, supplemented with 1 µM of all-trans-retinal. 16 h post transfection the cells were plated on poly-l-ornithine-coated 12-mm glass coverslips or 8-well µ-Slide (ibidi) at 20% confluence and analyzed by whole cell patch-clamp or confocal imaging 6-8 h later.

### Staining and live cell confocal imaging

HEK293T cells were stained with 5 µg/ml Hoechst 33342 to visualize nuclei and CellMask Deep Red (1:2000, Thermo Fisher Scientific) to label plasma membranes in Opti-MEM solution (Thermo Fisher Scientific). Imaging was performed at 37°C and 5% CO_2_ within 30 min after the staining on a Leica Stellaris 8 confocal microscope at the Microscopy Core Facility of the Medical Faculty at the University of Bonn. EYFP and CellMask Deep Red were excited with a tunable White Light Laser (440 – 790 nm) at wavelengths 514 nm and 653 nm, respectively. Hoechst 33342 was excited with a 405 nm laser. Middle-plane images were acquired using 63×/ 1.2 NA water immersion objective and additional ×3 zoom at 1024 × 1024 pixels and with line averaging of 8. The resulting images were processed in FIJI. *Er*NaR membrane localization was similar in at least 15 cells from three separate transfections.

### Patch-clamp recordings, data processing and statistics

Whole-cell patch clamp recordings were performed at room temperature, using a Multiclamp 700B amplifier (Molecular Devices). The signals were filtered at 10 kHz and digitized at sampling rate of 20 kHz with an Axon Digidata 1550B digitizer (Molecular Devices) using Clampex 11 Software (part of pCLAMP 11, Molecular Devices). Patch pipettes (3-6 MΩ) were fabricated from borosilicate glass with filament (GB150F-8P, Science Products GmbH) on a horizontal puller (Model P-1000, Sutter Instruments). Light was provided by a pE-800 system (CoolLED), controlled via TTL input, and connected to the optical path of Olympus SliceScope Pro 6000 upright microscope (Scientifica) via pE-Universal Collimator (CoolLED). LED light with maximum at 550 nm was applied at 34.3 mW/mm² irradiance in the focal plane of the 20×/ 0.5 NA water objective to activate *Er*NaR. To calculate the irradiance the output light power was measured using a bolometer (Coherent OP-2 VIS, Santa Clara) and divided by the illuminated area. The intensity distribution at the illuminated spot was fitted with gaussian and a radius, within which 95% of total power accumulates, was used as a measure of the spot size. The reference electrode was connected to the bath solution via an agar bridge with 150 mM KCl. The series resistance was <20 MΩ. Each cell was recorded three times and averaged directly in Clampex 11 Software to improve the signal-to-noise ratio. The ionic composition of the extracellular solution and all the intracellular solutions is indicated in Supplementary Table S3. Data were corrected for the respective liquid junction potential (LJP) after recording. LJPs for all solutions were measured directly and are stated in Supplementary Table S3. Data were analyzed using custom Wolfram Mathematica scripts and GraphPad Prism software. Photocurrent amplitudes were normalized to respective cell capacitance. Photocurrents at +60 mV were calculated from the linear fit of data points in positive voltages after LJPs correction. Time constant (t_off_) was determined by monoexponential fit of photocurrent decay upon light-off at holding voltage +80 mV. The data are presented as mean ± SEM of N = 6-9 cells for the photocurrent amplitudes or N = 5-8 cells for off-kinetics. The data from individual cells are also shown when appropriate. Photocurrents at +60 mV were tested for normal distribution using Shapiro-Wilk normality test (passed) and analyzed using two-way ANOVA with two Tukey’s multiple comparisons tests – for the effect of pH_i_ changes at fixed [Na^+^]_i_ and for the effect of [Na^+^]_i_ changes at fixed pH_i_. t_off_ was analyzed using Kruskal-Wallis test with Dunn’s multiple comparisons test.

### Protein expression, solubilization, and purification

*E.coli* cells were transformed with pET15b plasmid containing the gene of interest. Transformed cells were grown at 37°C in shaking baffled flasks in an autoinducing medium ZYP-5052^41^, containing 10 mg/L ampicillin. They were induced at an OD_600_ of 0.6-0.7 with 1 mM isopropyl-β-D-thiogalactopyranoside (IPTG). Subsequently, 10 μM all-*trans*-retinal was added. Incubation continued for 3 hours. The cells were collected by centrifugation at 5000g for 20 min. Collected cells were disrupted in an M-110P Lab Homogenizer (Microfluidics) at 25,000 p.s.i. in a buffer containing 20 mM Tris-HCl, pH 7.5, 5% glycerol, 0.5% Triton X-100 (Sigma-Aldrich) and 50 mg/L DNase I (Sigma-Aldrich). The membrane fraction of the cell lysate was isolated by ultracentrifugation at 125,000*g* for 1 h at 4°C. The pellet was resuspended in a buffer containing 50 mM NaH_2_PO_4_/Na_2_HPO_4_, pH 7.5, 0.15 M NaCl and 1% DDM (Anatrace, Affymetrix) and stirred overnight for solubilization. The insoluble fraction was removed by ultracentrifugation at 125,000*g* for 1h at 4 °C. The supernatant was loaded on a Ni-NTA column (Qiagen), and the protein was eluted in a buffer containing 50 mM NaH_2_PO_4_/Na_2_HPO_4_, pH 7.5, 0.15 M NaCl, 0.4 M imidazole, and 0.05% DDM. The eluate was subjected to size-exclusion chromatography on a Superdex 200i 300/10 (GE Healthcare Life Sciences) in a buffer containing 50 mM NaH_2_PO_4_/Na_2_HPO_4_, pH 8.0, 200 mM NaCl and 0.05% DDM. In the end, protein was concentrated to 70 mg/ml for crystallization and –80°C storage.

### Steady-state absorption spectroscopy and pH titration

Absorption spectra of *Er*NaR samples were measured with an absorption spectrometer (Specord600, Analytik Jena). Before and after each experiment, absorption spectra were taken to check sample quality. For the pH titration, samples were prepared to have a protein concentration of ∼0.4 mg/mL. The protein was suspended in the titration buffer containing 10 mM potassium citrate, 10 mM MES, 10 mM HEPES, 10 mM Tris, 10 mM CHES, 10 mM CAPS, and 100 mM Arg-HCl. The pH was adjusted with tiny amounts of [5000 mM] HCl or [5000 mM] KOH, respectively.

### Ultrafast transient absorption spectroscopy

Ultrafast transient absorption measurements were performed with a home-built pump-probe setup. A fs laser system - consisting of an Amplifier (Spitfire Ace-100F-1K, Spectra-Physics), seeded by a Ti:Sapphire oscillator (Mai Tai SP-NSI, Spectra-Physics) and pumped by a Nd:YLF laser (Empower 45, Spectra-Physics) - was used as the source for ultrashort laser pulses (100 fs, 800 nm, 1 kHz repetition rate). A home-built two-stage noncollinear optical parametric amplifier (NOPA) was used to generate the pump pulses at 550 nm. The white light continuum pulses used to probe the absorption changes of the sample were generated by focusing the 800 nm laser fundamental into a CaF_2_-crystal (3 mm). For the detection of the pump-probe signals a spectrometer (AvaSpec-ULS2048CL-EVO-RS, Avantes) was used. The measurements were performed in a 1 mm quartz cuvette and the sample was adjusted to have a protein concentration of ∼3.8 mg/mL protein. The sample was continuously moved in a plane perpendicular to the excitation beam to avoid photo-degradation. The excitation pulses were adjusted to an energy of 90 nJ/pulse.

### Transient flash photolysis spectroscopy

A Nd:YAG laser (SpitLight 600, Innolas Laser) was used to pump an optical parametric oscillator (preciScan, GWU-Lasertechnik). The OPO was set to generate excitation pulses with a central wavelength of 550 nm and an average pulse energy of ∼2.2 mJ/cm^2^. As probe light sources a Xenon or a Mercury-Xenon lamp (LC-8, Hamamatsu) were used. Two identical monochromators (1200 L/mm, 500 nm blaze), one in front and one after the sample, were used to set the chosen probing wavelengths. Absorption changes were detected by a photomultiplier tube (Photosensor H6780-02, Hamamatsu) and converted into an electrical signal afterwards. This signal was recorded by two oscilloscopes (PicoScope 5244B/D, Pico Technology) with overlapping timescales for detergent-solubilized samples and one oscilloscope (DPO5204B, Tektronix) for crystal samples. For each transient 30 acquisitions were measured and averaged to increase the S/N ratio. To obtain data files with a reasonable size for further analysis, raw data files were reduced using forward averaging and a combined linear and logarithmic timescale.

Detergent-solubilized samples were measured in a 2 x 10 mm quartz cuvette and prepared to have a concentration of ∼0.8 mg/mL protein. For conditions pH 8.0 0, 100, 1000 mM NaCl, pH 4.3 0, 100, 1000 mM NaCl, pH 4.3 1000 mM KCl and pH 4.3 1000 mM NMG, absorption changes were measured between 330 and 700 nm with a stepsize of 10 nm. Additional steps of the performed Na^+^ titration were measured by recording transients at characteristic wavelengths (pH 8.0: 340 nm, 450 nm, 540 nm, 610 nm and 620 nm; pH 4.3: 340 nm, 450 nm, 540 nm, 580 nm, 600 nm and 620 nm).

### Analysis of time-resolved spectroscopic data

Analysis of time-resolved spectroscopic data was performed using OPTIMUS software.^18^ The data of the ultrafast transient absorption and transient flash photolysis measurements were objected to the model-free lifetime distribution analysis (LDA) yielding the lifetime distributions of the individual photointermediate transitions, which are summarized in a lifetime density map (LDM).

### Cryo-EM grid preparation and data collection

For cryo-EM all samples were originally purified in the Buffer 1 (20 mM Tris pH 8.0, 200 mM NaCl, 0.05% DDM) and concentrated to 60 mg/ml using 100,000MWCO concentrator. For the structure of *Er*NaR at pH 8.0 (pH 8.0 dataset), the sample was diluted later with Buffer 1 (pH 8.0 structure). For the structure of *Er*NaR at pH 4.3 (pH 4.3 dataset), the sample was diluted with Buffer 2 (100 mM sodium acetate pH 4.6, 200 mM NaCl, 0.05% DDM) and run through Superdex200i 300/10 column in Buffer 2 to remove excess of Buffer 1. After that, the sample was again concentrated to 60 mg/ml 100,000MWCO concentrator. For grid preparation, all samples were diluted to 7 mg/ml, and volume of applied onto freshly glow-discharged (30s at 5 mA) Quantifoil grids (Au R1.2/1.3, 300 mesh) at 20 °C and 100% humidity and plunged-frozen in liquid ethane. The cryo-EM data were collected using 300 keV Krios microscope (Thermo Fisher), equipped with Gatan K3 detector.

### Cryo-EM data processing

All steps of data processing were performed using cryoSPARC v.4.0.2^42^ (Fig. S6). Motion correction and contrast transfer function (CTF) estimation were performed with default settings for all three datasets. Initial volume for template picking was generated after picking particles from pH 8.0 dataset using Topaz^43^ pre-trained model, followed by two rounds of 2D classification and *ab initio* model generation with 1 class, and homogeneous refinement. After that, for all datasets the final set of particles was picked using this volume as a template, with a template picker (150 Å particle radius), followed by duplicate removal with 50 Å distance.

For the pH 4.3 dataset, picked particles were extracted with 3x binning (384 px to 128 px). An initial set of particles was cleaned using two rounds of 2D classification (first round: 80 classes, 80 iterations, 5 final iterations, batch size 400, use clamp-solvent: true; second round: 40 classes, 40 iterations, batch size 200, use clamp-solvent: true). After that, particles were cleaned using a “3D classification” (*ab initio* model generation with 5 classes, followed by heterogeneous refinement), producing 555,294 particles for the non-binned refinement. These particles were re-extracted with 320 px box size, followed by homogeneous refinement (C5 symmetry, with per-particle CTF and defocus refinement) and local refinement (C5 symmetry, map generated with “Volume tools” at threshold 0.48), yielding a final resolution of 2.50 Å.

For the pH 8.0 dataset, picked particles were extracted with 4x binning (512 px to 128 px). An initial set of particles was cleaned using two rounds of 2D classification (first round: 50 classes, 40 iterations, batch size 200, use clamp-solvent: true; second round: 20 classes, 40 iterations, batch size 200, use clamp-solvent: true). After that, particles were cleaned using a “3D classification” (*ab initio* model generation with 5 classes, followed by heterogeneous refinement), producing 437,017 particles for the non-binned refinement. These particles were re-extracted with 640 px box size, followed by homogeneous refinement (C5 symmetry, with per-particle CTF and defocus refinement) and local refinement (C5 symmetry), yielding a final resolution of 2.63 Å.

### Model building and refinement

Automatically sharpened maps from cryoSPARC were aligned using UCSF ChimeraX^44^. The pentameric model of *Er*NaR was generated using Alphafold^45^ and docked as a rigid body into cryo-EM maps manually in ChimeraX. Further refinement was performed using Phenix^46,47^ and Coot^48^, producing the final statistics described in Table S1. Visualization and structure interpretation were carried out in UCSF Chimera^44,49^ and PyMol (Schrödinger, LLC).

### Crystallization

The crystals of *Er*NaR were grown with an *in meso* approach^50^, similar to that used in our previous works^12,19^. In particular, the solubilized protein (60 mg/ml) in the crystallization buffer was mixed with premelted at 42°C monoolein (MO, Nu-Chek Prep) in a 3:2 ratio (lipid:protein) to form a lipidic mesophase. The mesophase was homogenized in coupled syringes (Hamilton) by transferring the mesophase from one syringe to another until a homogeneous and gel-like material was formed.

Then, 150 nl drops of a protein–mesophase mixture were spotted on a 96-well LCP glass sandwich plate (Marienfeld) and overlaid with 400 nL of precipitant solution by means of the NT8 or Mosquito crystallization robots (Formulatrix and SPT Labtech, respectively). The best crystals were obtained with a protein concentration of 20 mg/ml (in the water part of the mesophase). The best crystals were obtained using 0.1M Sodium acetate pH 4.6, 10% PEG550MME (Hampton Research) as a precipitant. The crystals were grown at 22°C and appeared in 1 month.

For the determination of the *Er*NaR crystals structure at pH 4.6, once the crystals reached their final size, crystallization wells were opened, and drops containing the protein-mesophase mixture were covered with 100 μl of the precipitant solution. For the data collection, harvested crystals were incubated for 5 min in the precipitant solution. For the determination of the *Er*NaR crystals structure at pH 8.8, the crystals were originally grown at pH 4.6 using the same precipitant as described above. Once the crystals reached their final size, crystallization wells were opened and the crystals were soaked for 48 hours with exchanging the buffer three times. Crystals were harvested using micromounts (Mitegen, USA), flash-cooled, and stored in liquid nitrogen.

### Diffraction data collection and treatment

X-ray diffraction data of both structures of the ground state of *Er*NaR at pH 4.6 and 8.8 were collected at the P14 beamline of PETRAIII (Hamburg, Germany) using an EIGER X 16M and EIGER2 X 16M CdTe detectors. The data collection was performed using MxCube2 software. Diffraction images were processed using XDS^51^. The reflection intensities were scaled and merged using the Staraniso server^52^. There is no possibility of twinning for the crystals. In both cases, diffraction data from a single crystal were used. The data collection and treatment statistics are presented in Table S2.

### Crystal structure determination and refinement of *Er*NaR

Initial phases for the ground state of monomeric *Er*NaR at pH 4.6 were successfully obtained in the P6122 space group by molecular replacement using MOLREP^53^ from the CCP4 program suite^54^ using the 4XTL structure of monomeric KR2^9^ as a search model. The initial model was iteratively refined using REFMAC5^55^ and Coot^56^. The phases for the structure of *Er*NaR at pH 8.8 were determined using MOLREP with the phases of the *Er*NaR structure at pH 4.6 as a search model. The structure refinement statistics are presented in Table S2.

### Molecular simulation

Structures of all protonation models were optimized using electrostatic embedding QM/MM scheme^57,58^ using L-BFGS^59^ algorithm, implemented in Orca v5.0.3^60^ computational package. The QM part was treated with TD--DFT (B3LYP^61^ functional) level of theory, with Grimme^62^ dispersion correction, adopting the correlation-consistent^63^ cc-pVDZ atomic basis set and Resolution--of--Identity^64^ approximation. The MM part (part of protein within 5 Å of the retinal) was treated using the FF14SB AMBER force field^65^, while the rest of the protein was frozen. The vertical excitation energy was computed using the same QM/MM scheme at the RI second--order Algebraic Diagrammatic Construction (RI-ADC(2)) level of theory (QM part) implemented in the computational package Turbomole^66^. The point charges (MM part) of the environment were generated using the FF14SB AMBER force field.

## DATA AVAILABILITY

Atomic models built using X-ray crystallography and cryo-EM data have been deposited in the RCSB Protein Data Bank with PDB codes 8QR0 (for the pentameric form at pH 4.3), 8QQZ (for the pentameric form at pH 8.0), 8QLE (for the monomeric form at pH 4.6), and 8QLF (for the monomeric form at pH 8.8). Publicly available in RCSB Protein Data Bank structures of KR2 (PDB IDs: 4XTL, 6YC3, 6XYT) were used for analysis. The cryo-EM density maps have been deposited in the Electron Microscopy Data Bank under accession numbers EMD-18610 (pH 4.3) and EMD-18609 (pH 8.0).

### ACKNOWLEDGEMENTS

This research was supported by the German Research Foundation (CRC 1507 – Membrane-associated Protein Assemblies, Machineries and Supercomplexes; Project 05 to J.W.). G.H.U.L., A.V.S. and J.W. also thank Dr. Marvin Asido for helpful discussions. E.M., A.S. and A.G. thank NeCEN personnel for the support during data collection. The access to NeCEN facilities was funded by the Netherlands Electron Microscopy Infrastructure (NEMI), project number 184.034.014 of the National Roadmap for Large-Scale Research Infrastructure of the Dutch Research Council (NWO). A.G. was supported by NWO grant OCENW.KLEIN.141. We thank the Microscopy Core Facility of the Medical Faculty at the University of Bonn for providing support and instrumentation funded by the Bundesministerium für Bildung und Forschung (BMBF, Federal Ministry of Education and Research) – ACCENT: Förderung von Advanced Clinician Scientist im Bereich Immunopathogenese und Organdysfunktion, Gehirn und Neurodegeneration – Förderkennzeichen: 01EO2107. V.B. acknowledges support by the Volkswagen Foundation (Freigeist—A110720) and the Deutsche Forschungsgemeinschaft (EXC-2151-390873048-Cluster of Excellence— ImmunoSensation2 at the University of Bonn). K.K. has been supported by EMBL Interdisciplinary Postdoctoral Fellowship (EIPOD4) under Marie Sklodowska-Curie Actions Cofund (grant agreement number 847543). The work of A.A. was supported by funding from the German Research Foundation to T. Moser via the Multiscale Bioimaging - Cluster of Excellence (EXC 2067/1-390729940).

## AUTHOR CONTRIBUTIONS

K.K. performed bioinformatics analysis of the NDQ rhodopsins with the help of A.A. and under the supervision of A.B.; K.K. cloned the *Er*NaR gene for *E.coli* expression; C.B. and K.K. expressed and purified *Er*NaR in *E.coli*; T.B. supervised the expression and purification; E.P. cloned the *Er*NaR gene for electrophysiology studies with the help of N.M.; E.P. did the electrophysiology of *Er*NaR; G.H.U.L. and A.V.S. did the spectroscopy of *Er*NaR under the supervision of J.W.; K.K. crystallized the protein; R.A. helped with initial crystallization trials; V.G. supervised initial crystallization trials; K.K. collected and processed X-ray crystallography data and solved the crystal structures of *Er*NaR; G.B. helped with the X-ray crystallography data analysis; A.S. prepared grids and performed initial characterization and analysis of EM data; E.M. and A.G. processed cryo-EM data and obtained initial pentameric models of *Er*NaR; K.K. refined the cryo-EM structures; A.G. supervised grid preparation, cryo-EM data collection and processing and structure refinement and validation; D.F. performed QM/MM simulation under the supervision of I.S.; V.G. suggested the concept of short strong H-bonds for pKa control of functional residues; K.K., E.P., A.A., G.H.U.L., D.F., E.M. analyzed the data and prepared a manuscript with contributions of V.G., A.G., V.B., T.R.S., and all other authors.

## SUPPLEMENTARY INFORMATION

### Supplementary Figures

**Fig. S1.**
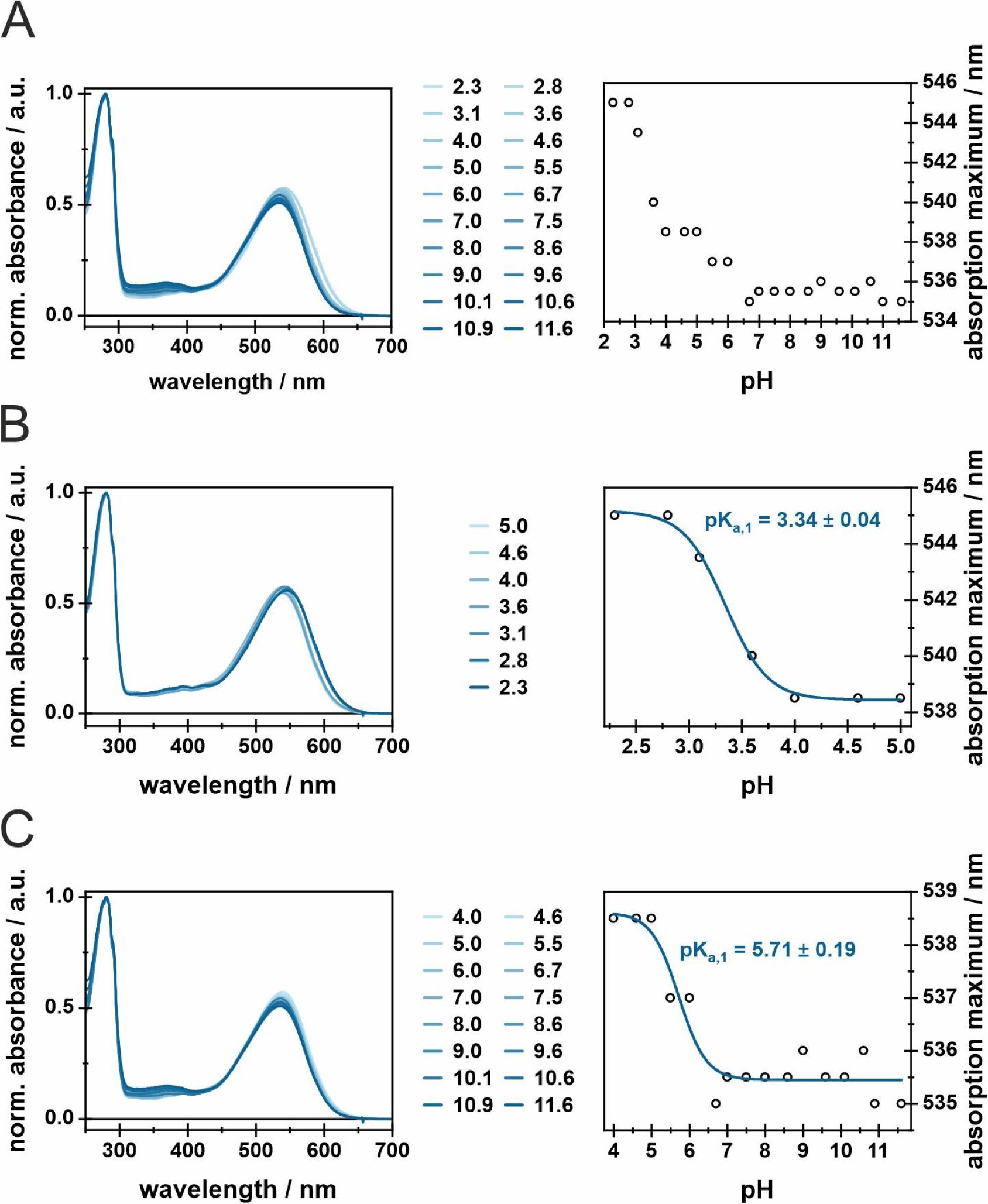
pH titration of detergent-solubilized *Er*NaR. Normalized absorption spectra of *Er*NaR for **A.** the complete investigated pH range (2.3 – 11.6), **B.** the pH range that corresponds to pK_a,1_(pH 2.3 – pH 5.0) and **C.** the pH range that corresponds to pK_a,2_ (pH 5.0 – 11.6). The spectra have been normalized to the absorbance at 280 nm. Lowering the pH value is accompanied by a spectral blue-shift of the main absorption band. In **B** and **C**, the observed trends were fitted with the Boltzmann function to obtain the pK_a_ values.

**Fig. S2.**
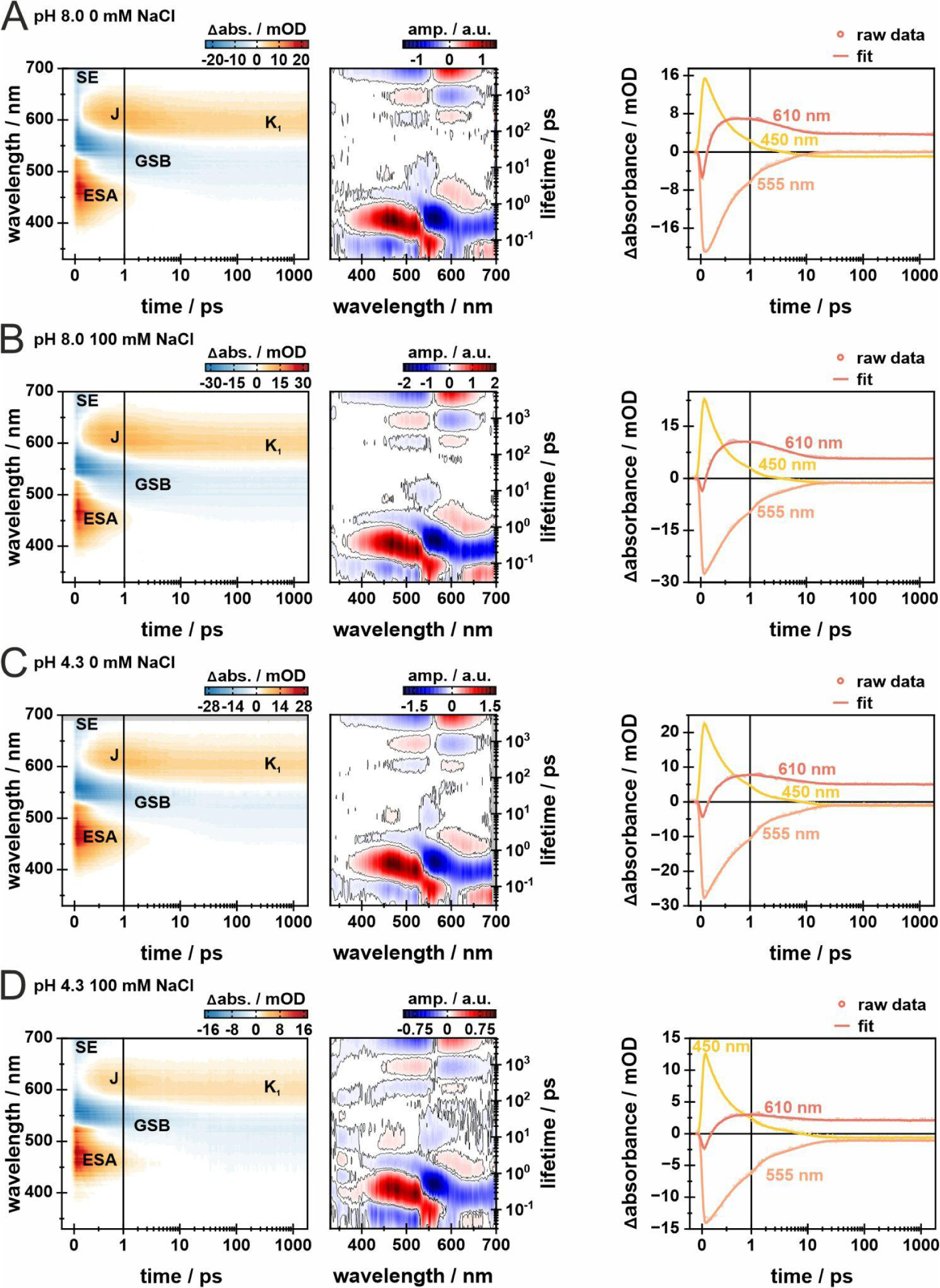
Ultrafast transient absorption data of *Er*NaR at different conditions. 2D-contour plots (left) and lifetime distribution maps (LDM) of ultrafast transient absorption measurements of *Er*NaR at **A.** pH 8.0 0 mM NaCl, **B.** pH 8.0 100 mM NaCl, **C.** pH 4.3 0 mM NaCl and **D.** pH 4.3 100 mM NaCl. Additionally, transients representative for the different signals at 450, 555 and 610 nm are shown as raw data (dots) and the obtained fit (lines) of the respective transients (right).

**Fig. S3.**
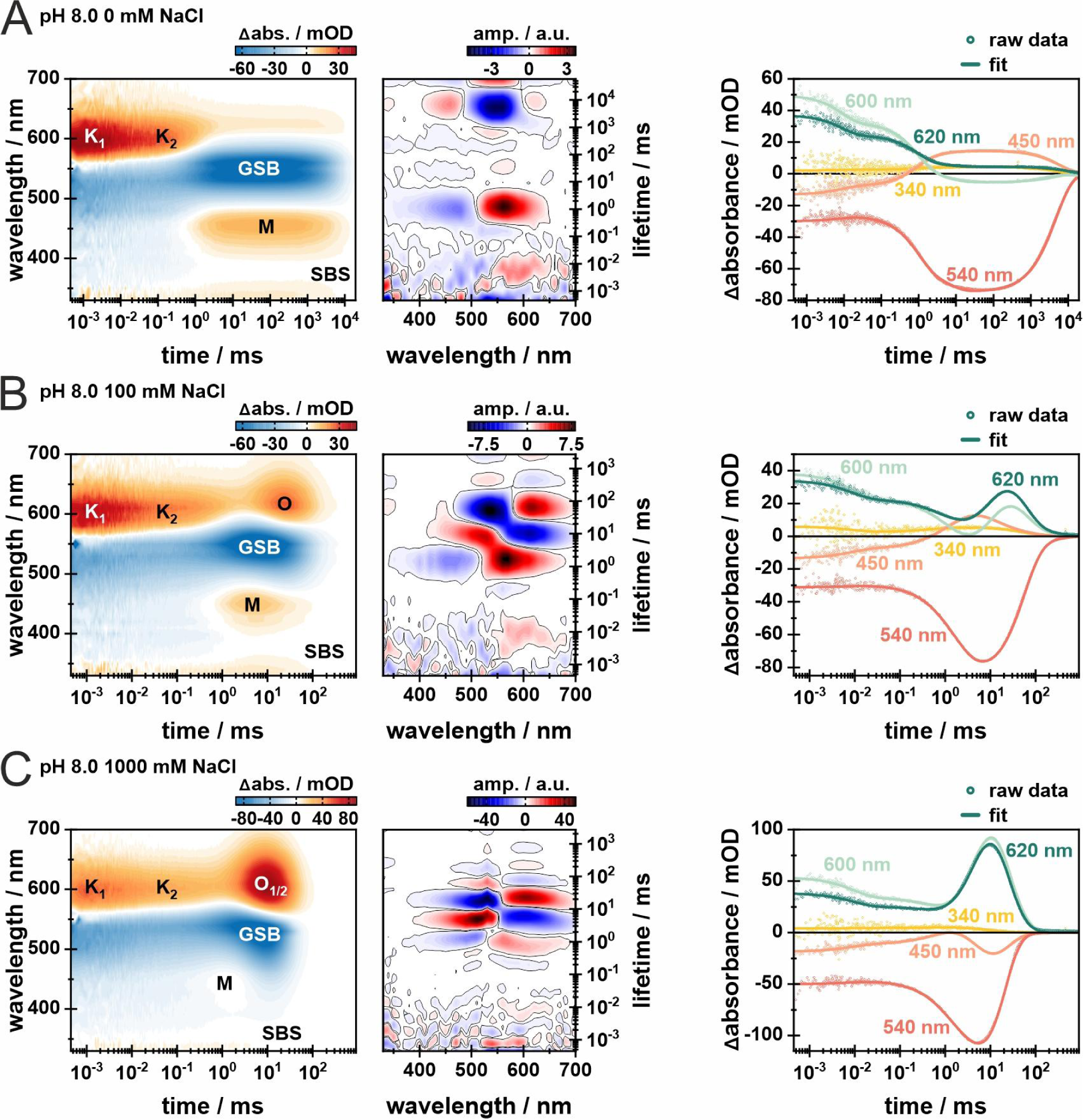
Transient flash photolysis data of *Er*NaR at pH 8.0 with different NaCl concentrations. 2D-contour plots (left) and lifetime distribution maps (LDM) of ultrafast transient absorption measurements of *Er*NaR with **A.** 0 mM NaCl, **B.** 100 mM NaCl and **C.** 1000 mM NaCl. Additionally, transients representative for the photocycle intermediates at 340 nm, 450 nm, 540 nm, 600 nm and 620 nm are shown as raw data (dots) and the obtained fit (lines) of the respective transients (right).

**Fig. S4.**
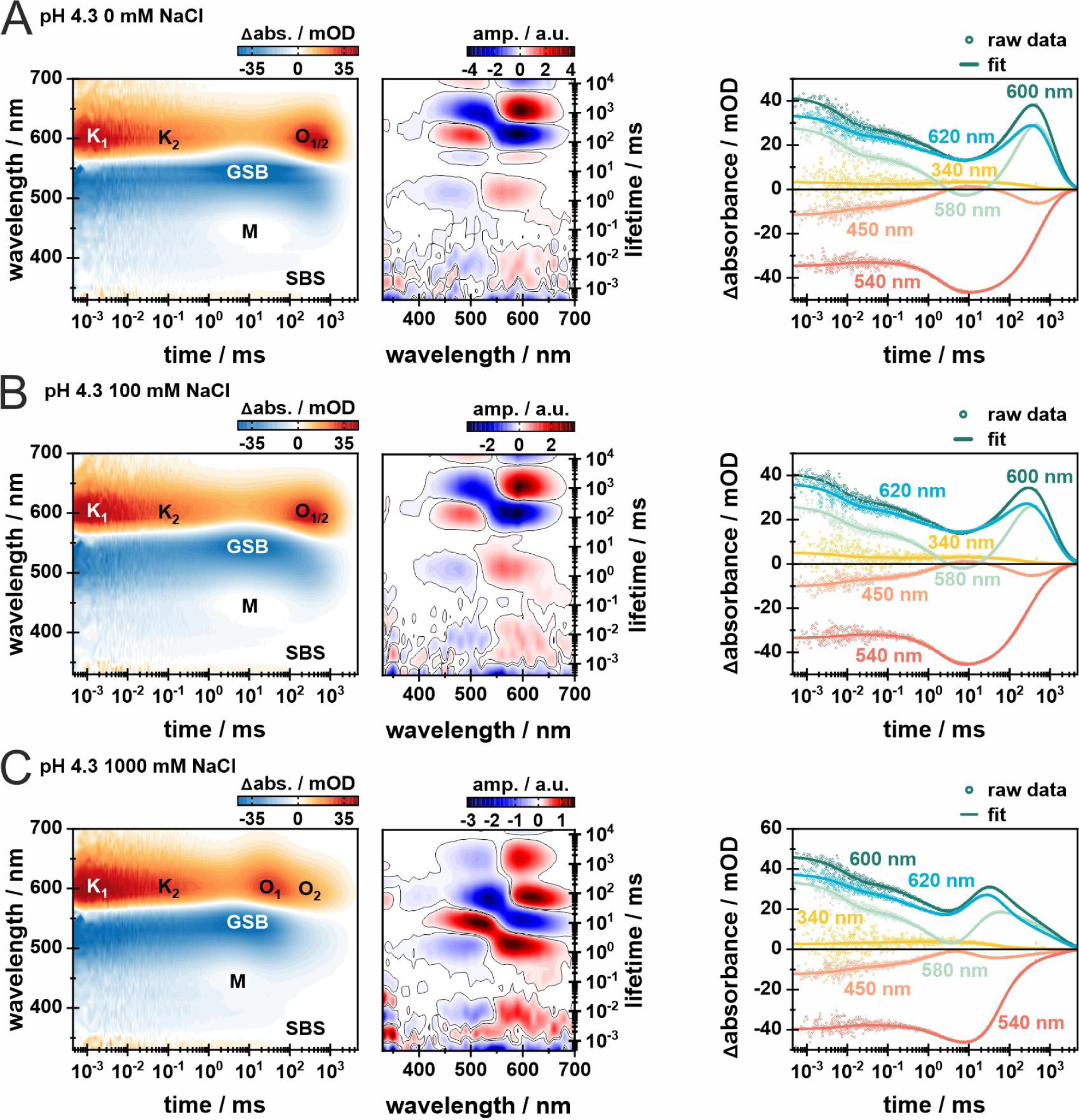
Transient flash photolysis data of *Er*NaR at pH 4.3 with different NaCl concentrations. 2D-contour plots (left) and lifetime distribution maps (LDM) of ultrafast transient absorption measurements of *Er*NaR with **A.** 0 mM NaCl, **B.** 100 mM NaCl and **C.** 1000 mM NaCl. Additionally, transients representative for the photocycle intermediates at 340 nm, 450 nm, 540 nm, 580 nm, 600 nm and 620 nm are shown as raw data (dots) and the obtained fit (lines) of the respective transients (right).

**Fig. S5.**
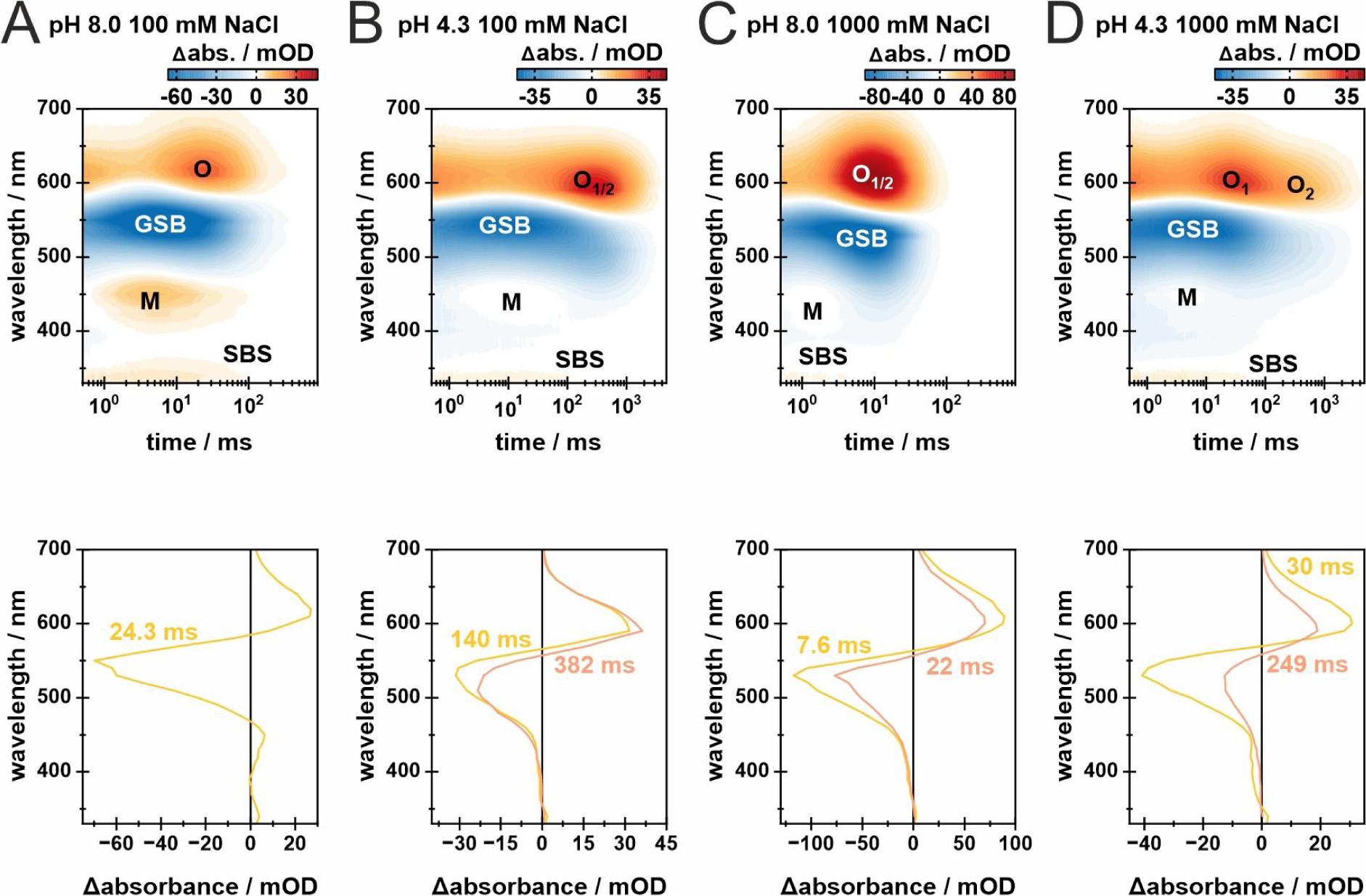
Sodium and pH dependence of the spectral position of the O_1_ and O_2_ states. 2D-contour plots as well as transient spectra at specific time points describe both O intermediates of the flash photolysis measurements of *Er*NaR at **A** pH 8.0 100 mM NaCl, **B** pH 4.3 100 mM NaCl, **C** pH 8.0 1000 mM NaCl and **D** pH 4.3 1000 NaCl. The 2D-contour plots are shown from 0.5 ms onwards till the end of the measurement to focus on the O_1_ and O_2_ states.

**Fig. S6.**
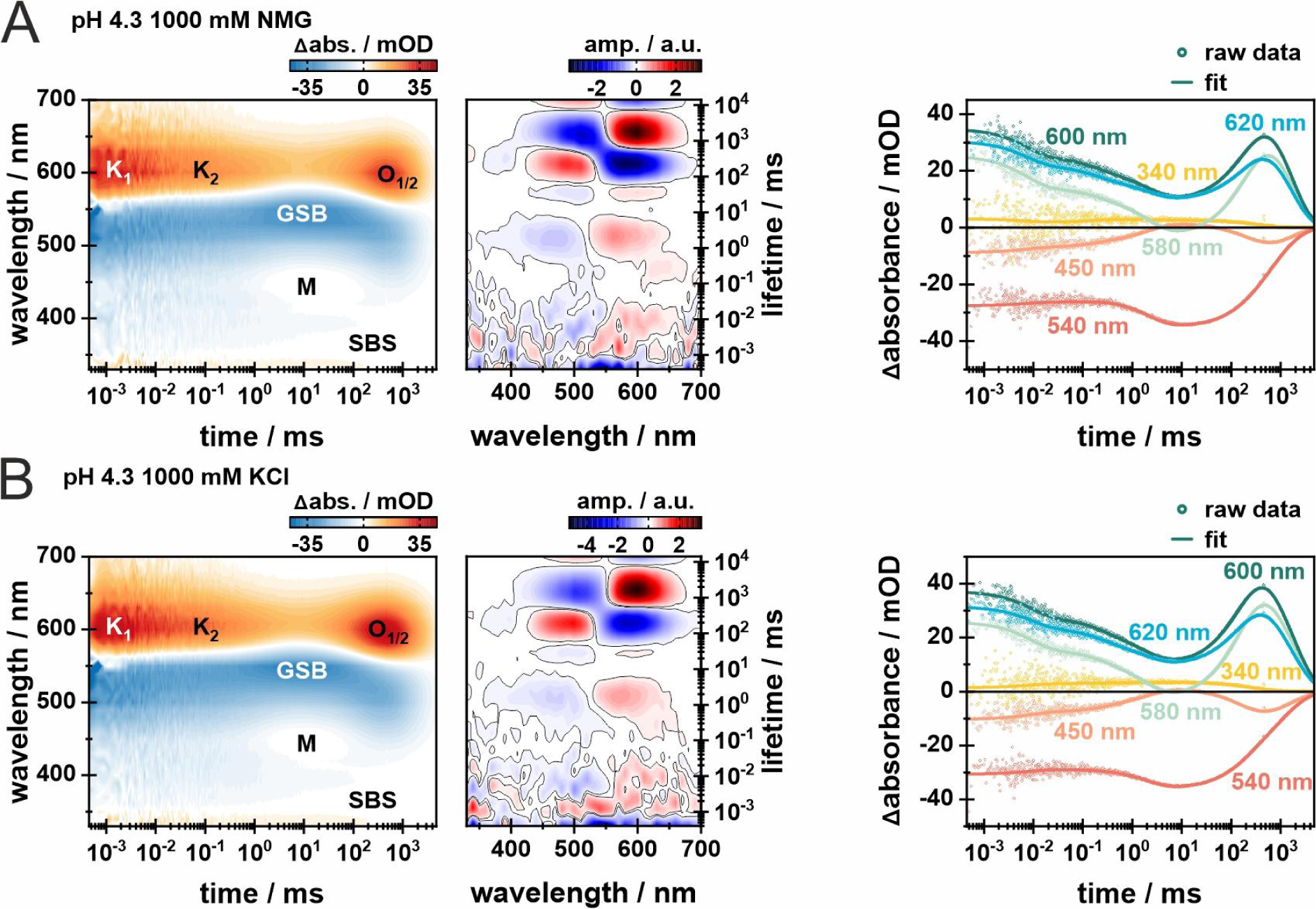
Transient flash photolysis data of *Er*NaR at pH 4.3 in the presence of different ions. 2D-contour plots (left) and lifetime distribution maps (LDM) of ultrafast transient absorption measurements of *Er*NaR with **A.** 1000 mM N-Methyl-D-glucamine (NMG) and **B.** 1000 mM KCl. Additionally, transients representative for the photocycle intermediates at **B** 340 nm, 450 nm, 540 nm, 600 nm and 620 nm and **B** 340 nm, 450 nm, 540 nm, 580 nm, 600 nm and 620 nm are shown as raw data (dots) and the obtained fit (lines) of the respective transients (right).

**Fig. S7.**
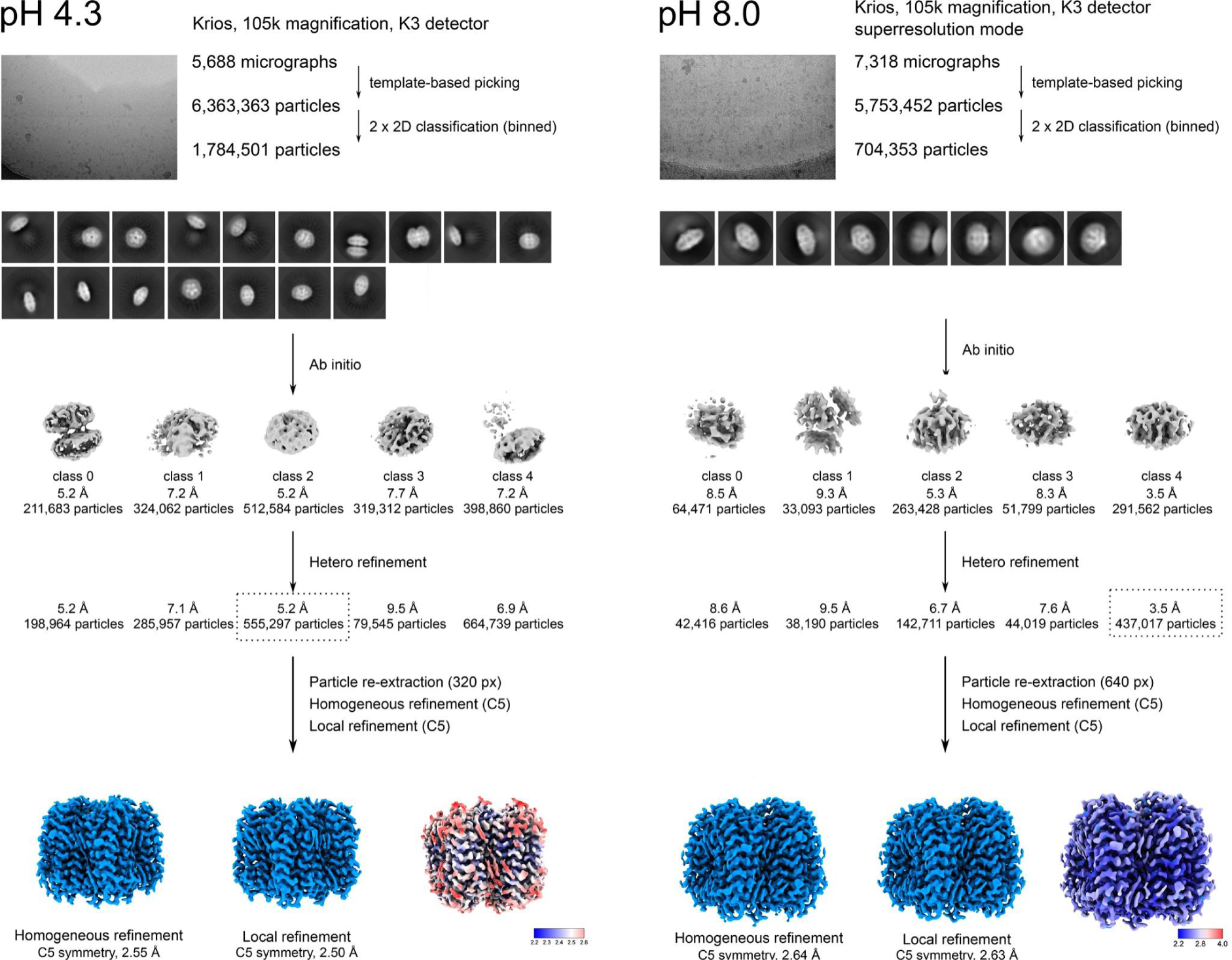
Workflow for solving the *Er*NaR structure using cryoSPARC. Overall resolution and local resolution are shown.

**Fig. S8.**
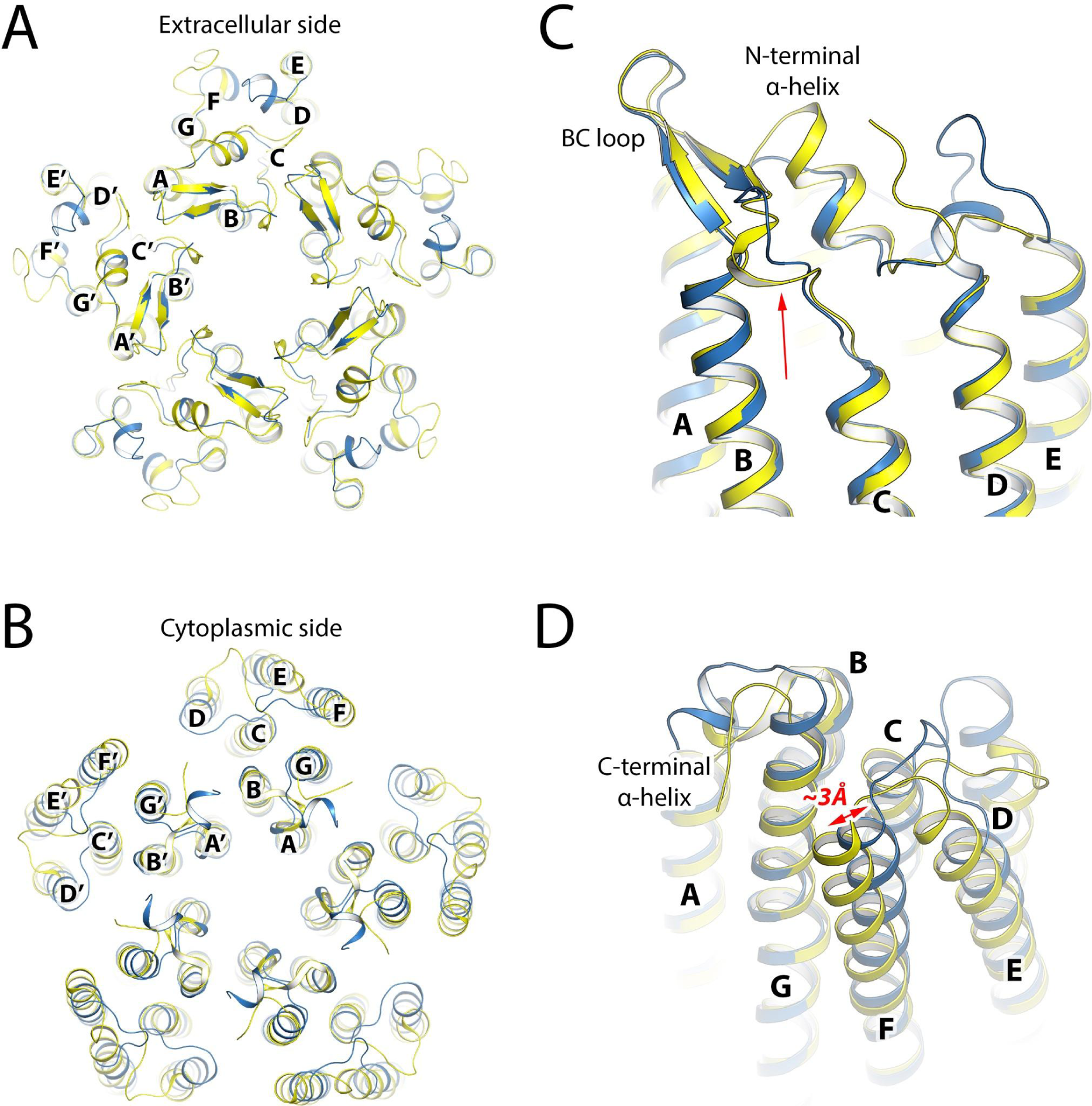
Comparison between the *Er*NaR and KR2 structures. **A.** Alignment of the pentamers of *Er*NaR (blue) and KR2 (yellow). View from the extracellular side. **B.** Alignment of the pentamers of *Er*NaR (blue) and KR2 (yellow). View from the cytoplasmic side. **C.** The extracellular side of the *Er*NaR and KR2 protomers. The red arrow indicates the end of the BC loop organized differently in KR2 and *Er*NaR. **D.** The cytoplasmic side of the *Er*NaR and KR2 protomers. The red arrow indicates the difference in the position of the cytoplasmic end of helix F in the *Er*NaR and KR2.

**Fig. S9.**
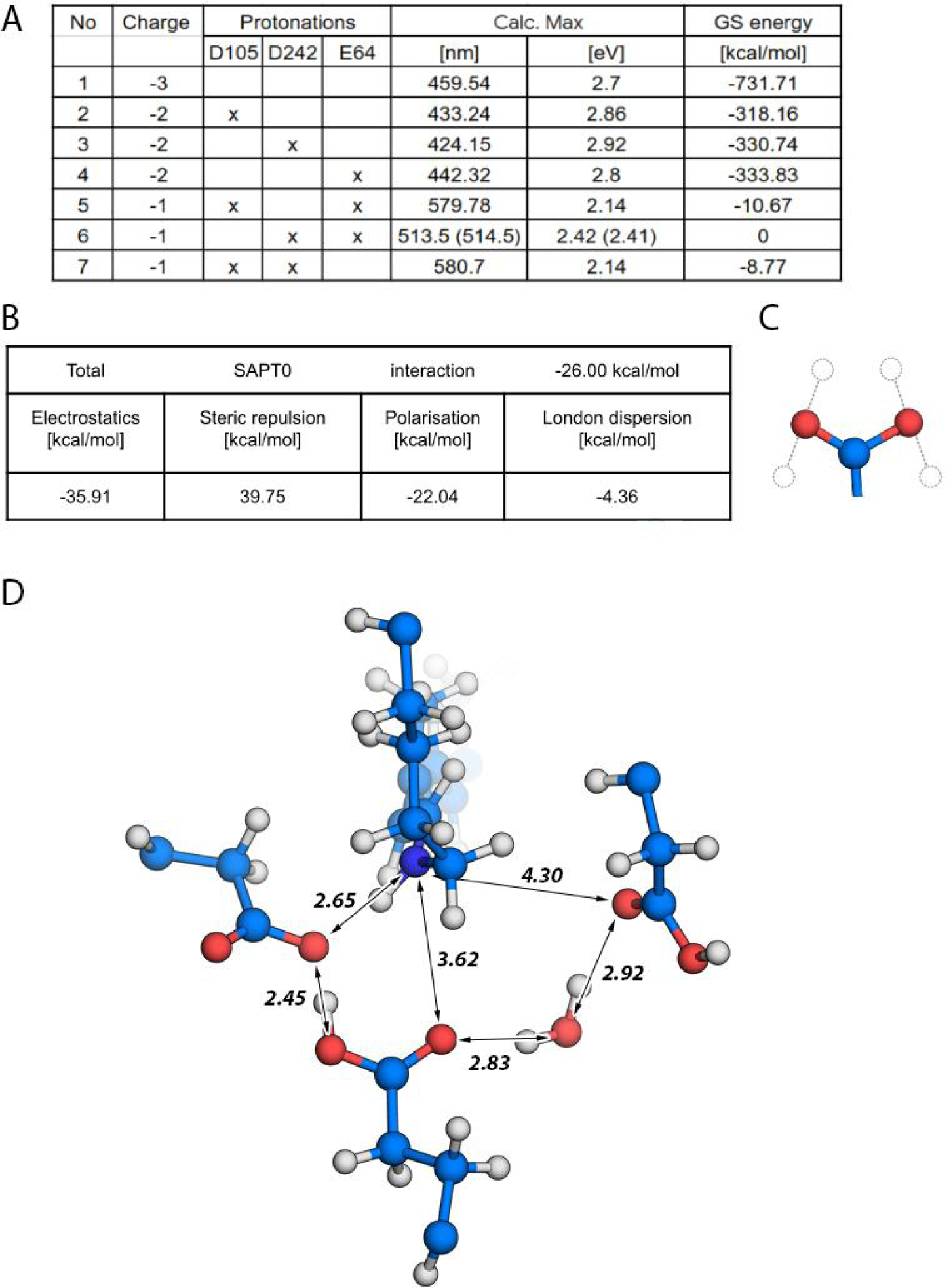
Overview of the QM/MM simulations. **A.** Various protonation patterns with the specification of individual protonation of counterions, excitation energies, and Ground State (GS) energies (values of GS energies were normalized with respect to the value of the lowest energy). Model 6 represents a protonation pattern the closest in agreement with the experiment (∼530 nm), as well as the lowest ground state energy compared to energies of other protonation systems. The wavelength in brackets corresponds to the inclusion of arginine (R98) for the quantum mechanics (QM) region, which slightly improves the agreement with the recorded spectrum. **B.** SAPT0 interaction energy components of D105 and E64 sidechains of model 6 of ErNaR. **C.** Orientations of the hydrogen atom for each of the three counter ions considered in QM/MM simulations. **D.** The visualization of model 6 and relative distances.

### Supplementary Tables

**Table S1.** Cryo-EM data collection, refinement, and validation.

|  | **4.3** | **8.0** |
| --- | --- | --- |
| **PDB ID** | 8QR0 | 8QQZ |
| **Data collection and processing** |  |  |
| Voltage (kV) | 300 | 300 |
| Electron exposure (e<sup>-</sup>/Å<sup>2</sup>) | 50.0 | 60.0 |
| Defocus range (μm) | −0.9 to −2.7 | −0.5 to −2.0 |
| Pixel size (Å) | 0.836 | 0.418 |
| Micrographs collected | 5,688 | 7,318 |
| Symmetry imposed | C5 | C5 |
| Initial dataset (# of particles) | 6,363,353 | 5,753,452 |
| Final dataset (# of particles) | 555,294 | 437,017 |
| Map resolution (Å) FSC<sub>01.43</sub> | 2.50 | 2.63 |
| **Refinement** |  |  |
| Initial model used | AlphaFold | AlphaFold |
| Map-sharpening *B* factor (Å<sup>2</sup>) | 111.4 | 111.0 |
| No. atoms |  |  |
| Protein | 10,735 | 10,801 |
| Retinal | 100 | 100 |
| Lipid | 618 | 775 |
| Water | 145 | 195 |
| *B*-factors (Å<sup>2</sup>) |  |  |
| Protein | 34.1 | 24.8 |
| Retinal | 23.6 | 13.5 |
| Lipid | 54.9 | 59.6 |
| Water | 41.0 | 30.6 |
| R.m.s. deviations |  |  |
| Bond lengths (Å) | 0.013 | 0.012 |
| Bond angles (°) | 1.787 | 1.633 |
| **Validation** |  |  |
| MolProbity score | 1.78 | 1.54 |
| Clash score | 5.64 | 4.98 |
| Poor rotamers (%) | 4.26 | 2.34 |
| Ramachandran plot (%) |  |  |
| Favored | 98.88 | 99.25 |
| Allowed | 1.12 | 0.75 |
| Outliers | 0.00 | 0.00 |
| Model to map fit CC | 0.92 | 0.90 |

**Table S2.**
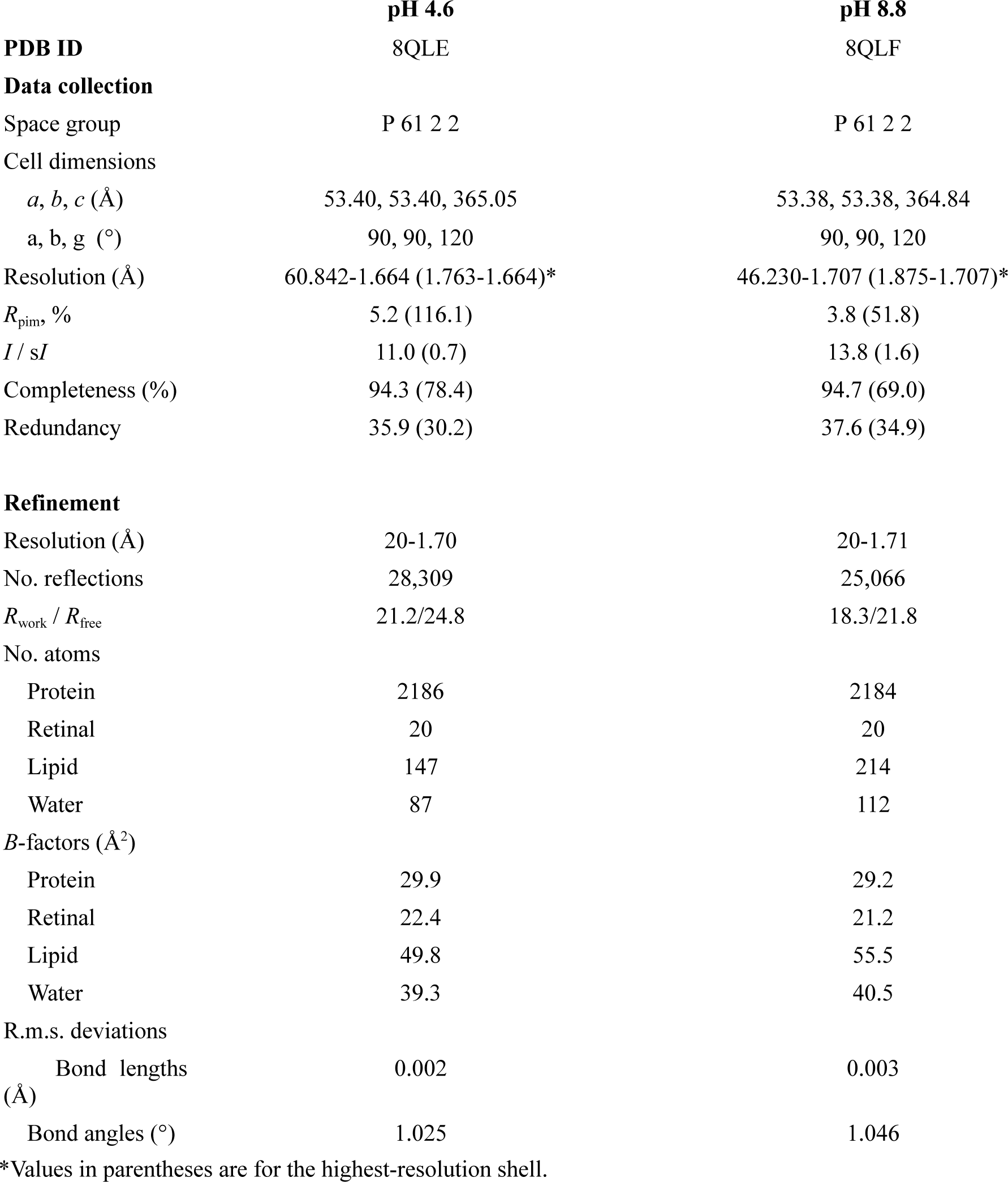
X-ray crystallography data collection and structure refinement statistics on *Er*NaR.

**Table S3.** Composition of extracellular and intracellular solutions and liquid junction potentials in whole cell patch-clamp experiments. All concentrations are given in mM. LJPs are in mV. The pH values of all solutions were very close to those of the buffers used but were adjusted if needed with 1M HCl or 1M KOH. Abbreviations: HEPES, 4-(2-hydroxyethyl)-1-piperazineethanesulfonic acid; EGTA, Ethyleneglycol-bis(β-aminoethyl)-N,N,Nʹ,Nʹ-tetraacetic Acid; Tris, tris(hydroxymethyl)aminomethane; LJP, liquid junction potential

|  | Extracellular solution | Intracellular solutions |  |  |  |  |  |  |  |  |
| --- | --- | --- | --- | --- | --- | --- | --- | --- | --- | --- |
|  |  | 130 mM Na<sup>+</sup> pH 7.5 | 20 mM Na<sup>+</sup> pH 7.5 | 0 mM Na<sup>+</sup> pH 7.5 | 130 mM Na<sup>+</sup> pH 5.0 | 20 mM Na<sup>+</sup> pH 5.0 | 0 mM Na<sup>+</sup> pH 5.0 | 130 mM Na<sup>+</sup> pH 9.0 | 20 mM Na<sup>+</sup> pH 9.0 | 0 mM Na<sup>+</sup> pH 9.0 |
| LJP (extracellular solution) | - | 0.5 | -7.3 | -7.9 | 0.6 | -5.3 | -6.6 | -0.4 | -7.5 | -8 |
| NaCl | 110 | 110 | - | - | 110 | - | - | 110 | - | - |
| Arginine-Cl | - | - | 110 | 110 | - | 110 | 110 | - | 110 | 110 |
| MgCl<sub>2</sub> | 2 | 2 | 2 | 2 | 2 | 2 | 2 | 2 | 2 | 2 |
| HEPES [NaOH] pH 7.5 | 10 | - | - | - | - | - | - | - | - | - |
| HEPES [KOH] pH 7.5 | - | 10 | 10 | 10 | - | - | - | - | - | - |
| EGTA [NaOH] pH 7.2 | - | 10<br>20 mM of Na<sup>+</sup> | 10<br>20 mM of Na<sup>+</sup> | - | 10<br>20 mM of Na<sup>+</sup> | 10<br>20 mM of Na<sup>+</sup> | - | 10<br>20 mM of Na<sup>+</sup> | 10<br>20 mM of Na<sup>+</sup> | - |
| EGTA [Tris] pH 7.2 | - | - | - | 10 | - | - | 10 | - | - | 10 |
| K-citrate buffer pH 5.0 | - | - | - | - | 10 | 10 | 10 | - | - | - |
| Tris buffer pH 9.0 | - | - | - | - | - | - | - | 10 | 10 | 10 |

